# Eukaryotic domestication of a bacterial immune protein following horizontal transfer

**DOI:** 10.64898/2026.04.30.722052

**Authors:** Edward M. Culbertson, Emily Cruz-Lorenzo, Jocelyn Leon Padilla, Megan Halfmann, James R. Drurey, Jeffrey J. Lange, Yao Li, Neha Garlapati, Harshitha Gompa, Benjamin R. Morehouse, Randal Halfmann, Tera C. Levin

## Abstract

Many components of eukaryotic innate immunity originated from bacterial immune systems. However, it is unclear how eukaryotes acquire these genes and how the components are domesticated into eukaryotic physiology. Here, we discovered a recent instance of bacteria-eukaryote horizontal transfer and used it to characterize the genetic and biochemical changes that accompanied eukaryotic acquisition. We focus on Toll/interleukin-1 receptor (TIR) domains, which are widespread yet potentially costly immune modules associated with inflammation and/or cell death. By generating an atlas of TIR diversity across the tree of life, we phylogenetically categorized the domains and uncovered highly diverged, eukaryotic TIR families. This analysis revealed a horizontal transfer event that created the TirBCD protein family of *Dictyostelium* amoebae, which is closely related to the bacterial immune protein TIR-STING. While the TIR domain was transferred into amoebae, the genomic locus did not contain known regulatory domains nor other components of a bacterial operon. Nevertheless, TirC retained biochemical and physiological similarities to TIR-STING. TirC is a highly potent NADase, capable of oligomerizing into large complexes and depleting NAD^+^ even at very low protein concentrations. When expressed in heterologous systems, TirC was spontaneously active and highly toxic. In contrast, the natural amoeba host tolerated expression of full length TirC, showing that the cells can typically regulate its activity and avoid autoimmunity. However, a truncated form of TirC induced rapid rounding and cell lysis. These results suggest that amoebae have used TirC to retool a form of bacterial cell death for use in eukaryotic cells. Overall, this study uncovers recent eukaryotic TIR evolution that captures features of both bacterial and eukaryotic immunity. We also expect that the TIR domain atlas will be useful to researchers across model systems as they explore the vast diversity of TIR molecular and cellular functions.

## Introduction

Recent studies have revealed that many proteins and domains used for innate immunity in animals are also shared with bacterial immunity, including Cyclic GMP-AMP synthase (cGAS), Stimulator of Interferon Genes (STING), argonaute, viperin, gasdermin, and Toll/interleukin-1 receptor (TIR) domains ^1–5^. We and others have found that these connections can arise through many evolutionary processes, including bacteria-eukaryote horizontal gene transfer (HGT) ^6–10^. However, many questions remain about how these gene transfers can be effective, as immune systems in bacteria and eukaryotes differ substantially in their genetic architectures, preferred immune proteins, and modes of innovation (i.e. HGT vs. gene duplication and divergence) ^11^. Additionally, because many of the proteins inhibit cell growth and/or trigger programmed cell death, acquisition of the proteins has a high potential to cause autoimmunity ^12^, particularly if they are misregulated following HGT. Thus, while we know these cross-kingdom gene transfers can occur, this process has been difficult to study as many of the immune gene acquisitions occurred long ago or in organisms that are not experimentally tractable. To better understand how eukaryotes acquire and co-opt bacterial proteins into their immune programs, we performed detailed evolutionary and experimental studies of TIR domains.

TIR domains are found in proteins throughout the tree of life, where they consistently play important roles in immunity ^13^. These domains have a conserved structure composed of a core of five parallel β-strands wrapped by five α-helices ^14^. Across animal, plant, and bacterial proteins, a common feature of TIR domains is that they require oligomerization to initiate immune responses ^15^. Bacteria and plants often use TIR domains to initiate the death or restrict the growth of infected cells via the enzymatic cleavage of β-nicotinamide adenine dinucleotide (NAD^+^). However, NAD^+^ processing can occur through different modalities. In *Sphingobacterium faecium* TIR-STING (*Sf*TIR-STING), a catalytic glutamate in the TIR domain is used to acutely deplete cellular NAD^+^, thus preventing bacteriophage replication and spread in the bacterial population ^7,16^. Alternatively, TIR domains can generate NAD^+^-based signaling molecules that activate downstream proteins, thus indirectly causing cell growth arrest or death of infected cells as in ThsB from the Thoeris antiphage defense system ^17^ and plant RPS4, a nucleotide-binding leucine-rich repeat (NLR) immune receptor ^18^. NAD^+^ hydrolysis by enzymatic TIR domains often generates nicotinamide (NAM) and ADP-Ribose (ADPR) as byproducts, although numerous other linear and cyclic ADPR isomers are also possible ^19^. Within animals, TIR domains are most commonly found in Toll-like receptors and related proteins and adaptors ^20^. These TIRs are primarily thought to function as scaffolds, recruiting other proteins via TIR-TIR oligomerization to build large signaling complexes that initiate inflammatory regulation ^21^. Later work discovered enzymatic TIR proteins in animals such as the sterile alpha and TIR motif-containing protein-1 (SARM1) protein ^22^, which depletes NAD^+^ in neurons resulting in neuronal cell death upon injury ^23^. Interestingly, phylogenetic analyses suggested that the TIR domain in SARM1 is closely related to bacterial TIR domains and was likely acquired through HGT ^24^. While our understanding of TIR-mediated immunity in bacteria, plants, and animals is broadening, expanded analyses of bacterial TIRs continue to find additional mechanisms of TIR-mediated immunity. We expect the same will be true when we sample eukaryotes more thoroughly.

Amoebae are microeukaryotes found in soil and aquatic environments, where they feed on environmental bacteria ^25^. Efficient bacterial predation and evasion of intracellular bacterial infections requires amoeba cells to use robust antibacterial mechanisms such as phagosome-lysosome fusion, nutritional immunity, generation of bactericidal reactive oxygen species, generation of extracellular DNA traps, and autophagy ^26^. Some of these responses have been linked to the TIR-containing *Dictyostelium* protein TirA ^27^ ^28^, although the molecular and cellular roles of TIR domains in amoeba immunity are not well understood. While bacteria, plants, and animal SARM1 use TIR proteins to regulate programmed cell death, there is currently no evidence that this occurs in microeukaryotes. While *Dictyostelium discoideum* does undergo a caspase-independent, vacuolar programmed cell death under certain developmental or physiological conditions ^29–31^, there are no known roles for TIR-containing proteins in this process.

To understand how amoeba TIR domains are related to TIRs in other species, we used phylogenetics to categorize the relationships among thousands of TIR domains across bacteria, archaea, and diverse eukaryotes. We generated a TIR domain atlas that allowed us to create new phylogenetic designations for TIRs, with a focus on eukaryotic TIRs. In the process, we discovered homologs of bacterial immune genes that were recently horizontally transferred to *Dictyostelium* amoebae. The TIR domains from *Dictyostelium* TirBCD proteins are closely related to the TIR from *Sf*TIR-STING, although they do not share regulatory domains. Through biochemical, cellular, and genetic experiments, we found that amoeba TirC is a highly active NADase that oligomerizes into filaments and causes toxicity when expressed in *E. coli* or yeast cells. Expression of full length TirC was tolerated within amoeba cells, indicating that it has been successfully domesticated by this new host. However, when expressing TirC without its N-terminal transmembrane domain, we observed toxicity and cell lysis, suggesting that the co-opted protein may be involved in programmed cell death analogous to processes in bacterial, plant, and animal immunity. Overall, this system can serve as a useful model for how eukaryotes adapt and deploy recently-acquired bacterial immune genes.

## Results

### An atlas of TIR domains across eukaryotes

Due to the key roles of TIR domains in immune defenses across many species, we sought to characterize the diversity of eukaryotic TIR proteins and understand their relationships to prokaryotic TIRs. As TIRs are abundant and highly diverse domains, they have previously been categorized into six Pfam categories within the STIR clan, which includes the SEFIR domains as an outgroup to TIRs ^32^. We began by gathering sequences through iterative hidden Markov model (HMM) searches of the EukProt v3 database of nearly 1000 diverse eukaryotic species, incorporating the hits from each search into the following HMM ^6^. This iterative process was repeated for all six of the published PFAM HMMs comprising the STIR clan, searching each HMM iteratively until saturation before adding on the next PFAM group (Supp. Fig 1). Ultimately, we collected 7,432 TIR domains from across eukaryotes. We compared the eukaryotic sequences to 1,196 representative bacterial and archaeal TIRs from each PFAM category. We then downsampled this sequence set to obtain a phylogenetically diverse set of 4,105 TIRs (3,082 eukaryotic + 810 bacterial + 120 archaeal). We aligned the TIR domains, trimmed poorly aligned residues from the alignment, and used IQTree to generate a TIR phylogeny. We generated four such trees, made from different alignment algorithms and different trimming criteria and then compared which clades remained coherent across the trees (Supp. Fig. 2). We thus identified 13 supported TIR subclades in addition to the SEFIR domain outgroup (Fig. 1A & B, Supp. Table 1).

**Figure 1.**
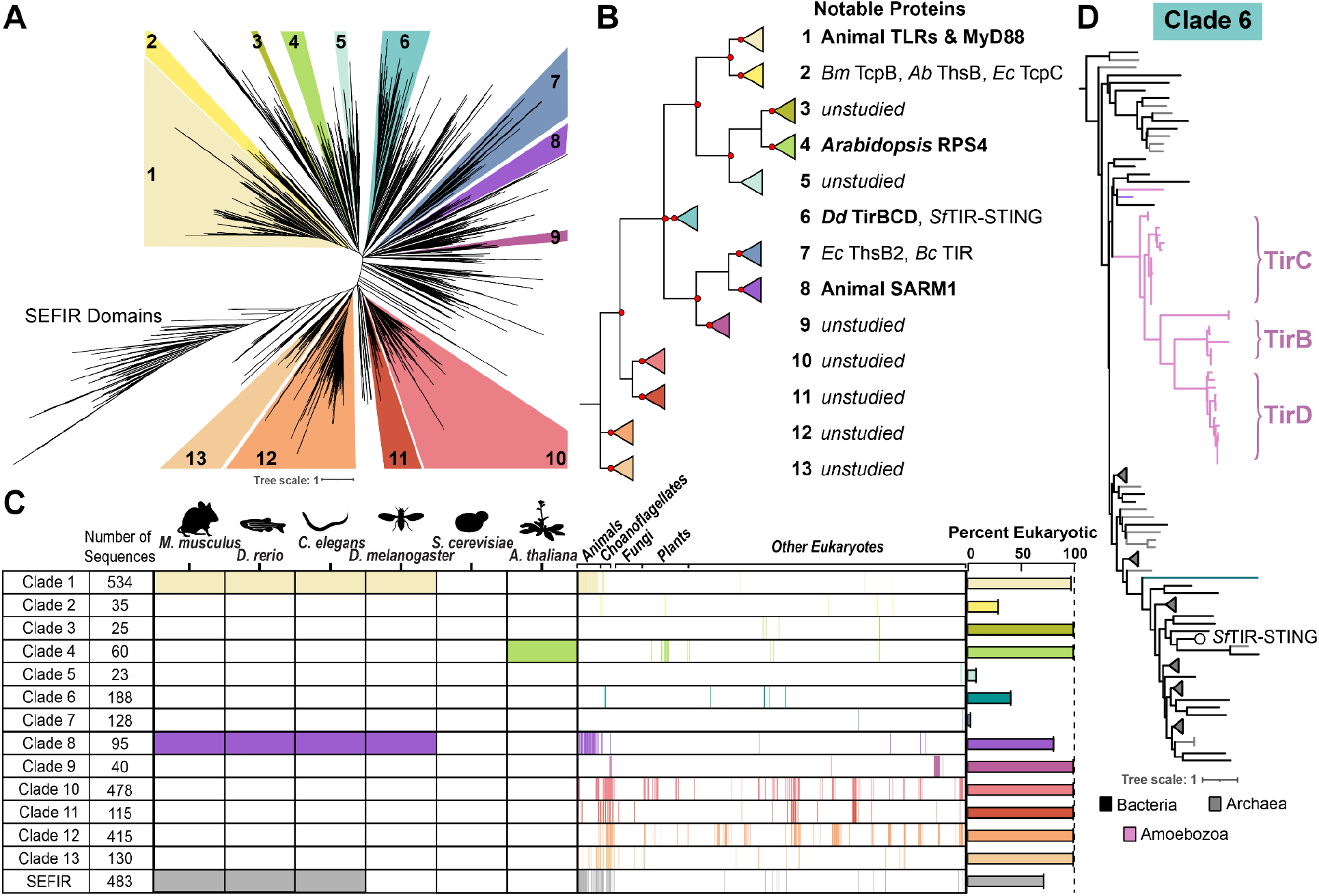
Atlas of diverse eukaryotic TIR domains identifies recently horizontally acquired amoeba TIR proteins. **A.** Maximum likelihood phylogenetic tree generated by IQtree of TIR domains from bacteria, archaea, and eukaryotes. Numbered, colored clades represent portions of the tree that contain at least 20 sequences and are supported by ultrafast bootstraps >70 in multiple IQtrees built from different alignments (Supp. Fig 2). TIR branches that lie outside of colored clades had variable placements across different trees. **B.** A topological overview of the phylogenetic tree in (A), red dots indicate nodes with >50 ultrafast bootstraps. A selection of previously studied proteins found in each clade are listed. *Bm* = *Brucella melitensis*, *Ab* = *Acinetobacter baumannii*, *Ec* = *E. coli*, *Sf* = *Sphingobacterium faecium*, *Bc* = *Bacillus cereus*. **C.** Prevalence of different TIR domains across eukaryotes. Commonly studied model organisms encode TIRs from clades 1, 4, and/or 8, whereas non-model eukaryotes encode clades 1-13. For eukaryotic species, the presence of 1+ TIR from that clade is represented by a colored box. On the right is a bar chart detailing the percentage of sequences in each clade of the phylogeny that are from eukaryotic organisms. **D.** A maximum likelihood phylogenetic tree generated by IQtree of Clade 6, illustrating where amoeba TirBCD proteins (pink) branch relative to bacterial (black) and archaeal (gray) CAP12/Pycsar TIR domains. All nodes connecting amoeba and bacterial sequences are supported by ultrafast bootstraps >85.

**Figure 2.**
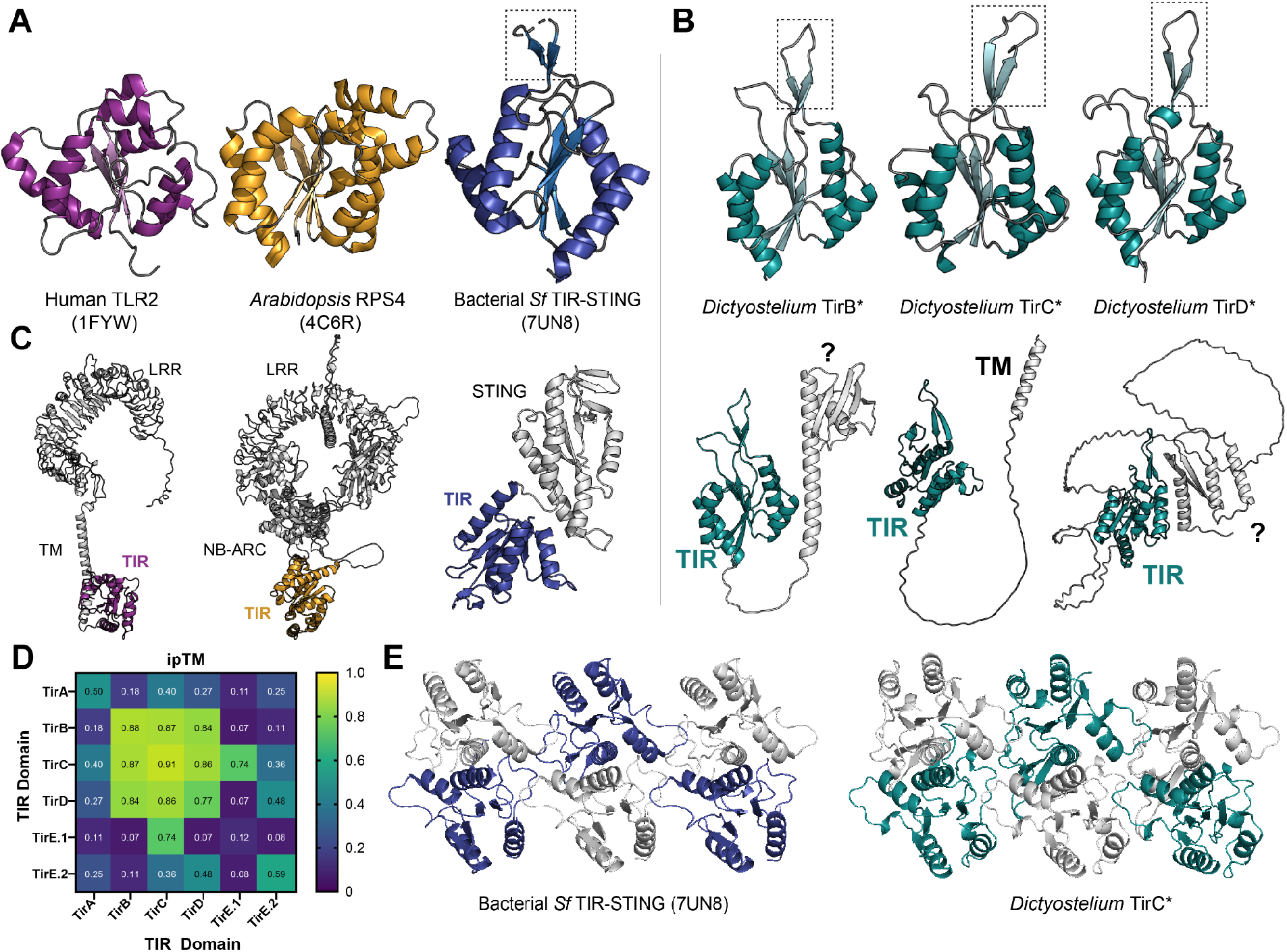
Amoeba TirC is predicted to homo-oligomerize similar to *Sf*TIR-STING. **A.** Crystal and cryo-EM structures of the TIR domains of human TLR2 (PDB: 1FYW), *Arabidopsis* RPS4 (4C6R), and *Sf*TIR-STING (7UN8). **B.** AlphaFold predictions of the TIR domains of *Dictyostelium discoideum* TirB, C, and D. The dotted box outlines the C-C loop extension. **C.** AlphaFold predictions of the full length proteins shown in A and B. Other domains are labeled: leucine-rich repeats (LRR), transmembrane domain (TM), NB-ARC domain, STING domain, and TIR domains. The N-terminal domains in TirB and TirD proteins are unidentified (?). TirC contains a hydrophobic N-terminal helix that could be a transmembrane domain. **D.** Dimerization potential across all of the TIR domain proteins in *Dictyostelium*, predicted by AlphaFold ipTM scores. Because TirE encodes two TIR domains, pairwise dimerization scores are shown for each domain separately. **E.** CryoEM model of the TIR domains of the *Sf*TIR-STING filament (left) and AlphaFold model of a TirC hexamer (right). In both, alternating TIR domains are colored in blue or teal to highlight the similarity of the hexamer structures.

To determine how our clade designations compared to the prior TIR categories, we labeled each sequence with its top PFAM hit and assessed where these annotations were found across the trees. The TIR_2 Pfam HMM (PF13676) hit an extremely broad set of TIR domains, including sequences from across the tree in nearly every subclade. Therefore, TIR_2 is useful for defining a sequence as a TIR domain, but not for categorizing subtypes of TIRs. In contrast, TIR (PF01582) corresponded largely to clade 1. CAP/Pycsar TIRs (PF10137) were exclusively localized to clade 6. ThsB-TIR (PF08937) was found predominantly in clades 5 and 7, and all of the clade 7 TIRs corresponded to this PFAM domain. TIR_3 (PF18567) preferentially hit a small collection of metazoan TIRs containing BANK1 and PIk3ap1/BCAP. This cluster was not well supported and did not meet our requirements to assign it into a reproducible clade. Nevertheless, we believe the TIR clades defined here are useful, because they identify reliable subgroups of sequences while largely retaining the phylogenetic structure of prior categorizations.

We then asked where previously studied TIR proteins were found in the trees. Clade 1 was the largest group of TIRs and consisted almost entirely of animal sequences, including those from all Toll-like receptors, interleukin receptors, the TLR adaptor protein Myeloid differentiation primary response 88 (MyD88), and Toll-interleukin-1 Receptor domain-containing adaptor protein (TIRAP) (Fig. 1B, Supp. Fig 3). All of the TIR domains from land plants (Embryophyta) fell into Clade 4. For both the Clade 1 animal and Clade 4 plant sequences, most of the TIRs within the clades were closely related, reflecting a long history of repeated immune gene duplication and diversification that has occurred independently in animals and plants ^33^. While we were primarily focused on eukaryotic TIRs, we also investigated the placement of bacterial and archaeal sequences on the trees (although see ^34^ and prior work for a specific focus on bacterial TIR diversity). Most bacterial sequences were found within clades 2, 5, 6, and 7. Clade 7 contained a mix of bacterial and archaeal TIRs, including ThsB TIRs from *E. coli* and *Bacillus cereus.* ^17,35^. Clade 2 contained both *Acinetobacter baumannii* ThsB and *E. coli* TcpC, despite the fact that these two TIRs process NAD^+^ into different ADPR isomers ^36^. Therefore, TIR phylogenetic similarity does not necessarily equate to similar biochemical function. Finally, Clade 8 included the TIR from animal SARM1, which plays important roles in neuronal cell death through its NADase activity ^23,37,38^. Our results support previous findings that SARM1’s TIR domain arose through horizontal transfer from bacteria ^24^, as SARM1 branches robustly within a bacterial lineage and its Clade 8 location is distinct from all other animal TIRs (Supp. Fig. 3). In addition, we identified a small number of SARM1-like sequences from choanoflagellates, amoebae, green algae, and dinoflagellates, suggesting that this may represent an ancient horizontal acquisition, possibly followed by additional transfers within eukaryotes.

**Figure 3.**
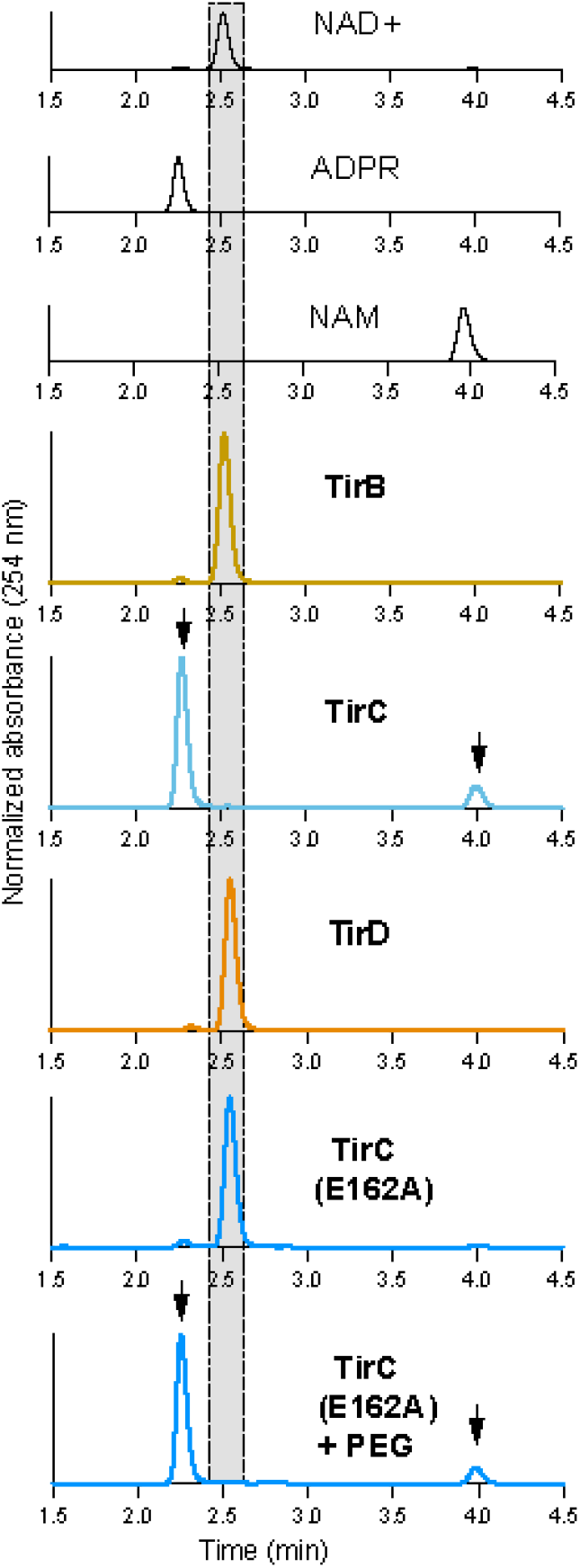
TirC is an active NADase. High-performance liquid chromatography analysis of NAD^+^ incubated with purified TirB, TirC, TirD, and the TirC E162A mutant. Shaded box highlights the unprocessed NAD^+^ peak. Black curves show standards while colored curves show the results of co-incubations with amoeba proteins. No NAD^+^ processing was observed following co-incubations with TirB or TirD. In contrast, TirC cleaves NAD^+^ into ADP-ribose (ADPR) and nicotinamide (NAM), indicated by arrows. The TirC E162A protein was inactive following mutation of the predicted catalytic glutamate, except in the presence of 10% PEG 8000 as a molecular crowding agent. Normalization was done within each sample to set the maximum peak height value to 100% and baseline to 0%.

Beyond these characterized TIR proteins, eukaryotic sequences were distributed across the tree. Indeed, eukaryotic TIRs in our dataset made up the entirety of clades 3, 9, 10, 11, 12, and 13, none of which have been characterized to date. Notably, clades 10-13 contain TIR domains that are highly diverged from any previously studied proteins, yet are comparatively common across eukaryotes, including an array of animal species (e.g. non-model Arthropods, Molluscs, and Cnidarians) (Fig. 1B-C, Supp. Table 1). These sequences have likely been overlooked to date because routinely studied eukaryotic model organisms encode TIRs from only clades 1, 4, and 8 (Fig. 1C). However, given the diversity of enzymatic activities and immune roles already discovered within characterized TIR domains ^14,39–41^, we expect that these unstudied TIRs represent potential new modalities of TIR function across diverse eukaryotes.

### Amoeba TIR domains were horizontally acquired from anti-phage immune proteins

In Clade 6 we discovered an unusual instance of TIR domain bacteria-eukaryote horizontal transfer that has occurred quite recently (Fig. 1D). The amoeba *Dictyostelium discoideum* encodes five TIR domain proteins: TirA, TirB, TirC, and two proteins that we identified here and named TirD and TirE. Previously designated as *DDB_G0278125*, TirE contained multiple predicted RCC1 repeats as well as two diverged, tandem TIR domains that were recognized by our expanded TIR sampling but not by prior PFAM HMMs. The *tirD* locus was previously designated as two genes, *DDB_G0287321* & *DDB_G0287437*, due to the presence of a stop codon separating the two genes in the original reference genome ^42^. However, other recently sequenced *D. discoideum* genome sequences lack this stop codon and we experimentally verified that the locus encodes a single in-frame gene (Supp. Fig. 4C, ^43^). We found that TirB, TirC, and TirD are paralogs that all branch with strong statistical support within a group of CAP12/Pycsar family bacterial and archaeal TIR sequences (Fig. 1D). Several features indicated that the TirBCD sequences have been domesticated following HGT and are now bona fide amoeba proteins in *D. discoideum*. First, the TirBCD sequences formed coherent clades in the phylogeny, consistent with vertical inheritance across amoeba species (Fig 1D). Second, the genes had a low GC content (19-28%) and TirC/TirD had poly-N tracts, features common to *Dictyostelium* genes ^42–44^. Third, the *tirB, tirC,* and *tirD* genes were each found in consistent syntenic loci, demonstrating that they reside within amoeba genomes and do not arise from bacterial contamination (Supp. Fig 5). Fourth, the *tirB* and *tirD* gene models contained predicted introns, which we validated using PCR from amoeba cDNA (Supp. Fig. 4A & C). Finally, these paralogs have distinct patterns of gene expression in amoebae: *tirB* is upregulated during multicellular development, while *tirC* is upregulated during hypoxia ^45,46^ (Supp. Fig. 4E & F). Thus, we conclude that the TIR domains of TirBCD were horizontally transferred from prokaryotes into amoebae, where they have since been incorporated into amoeba cell biology.

**Figure 4.**
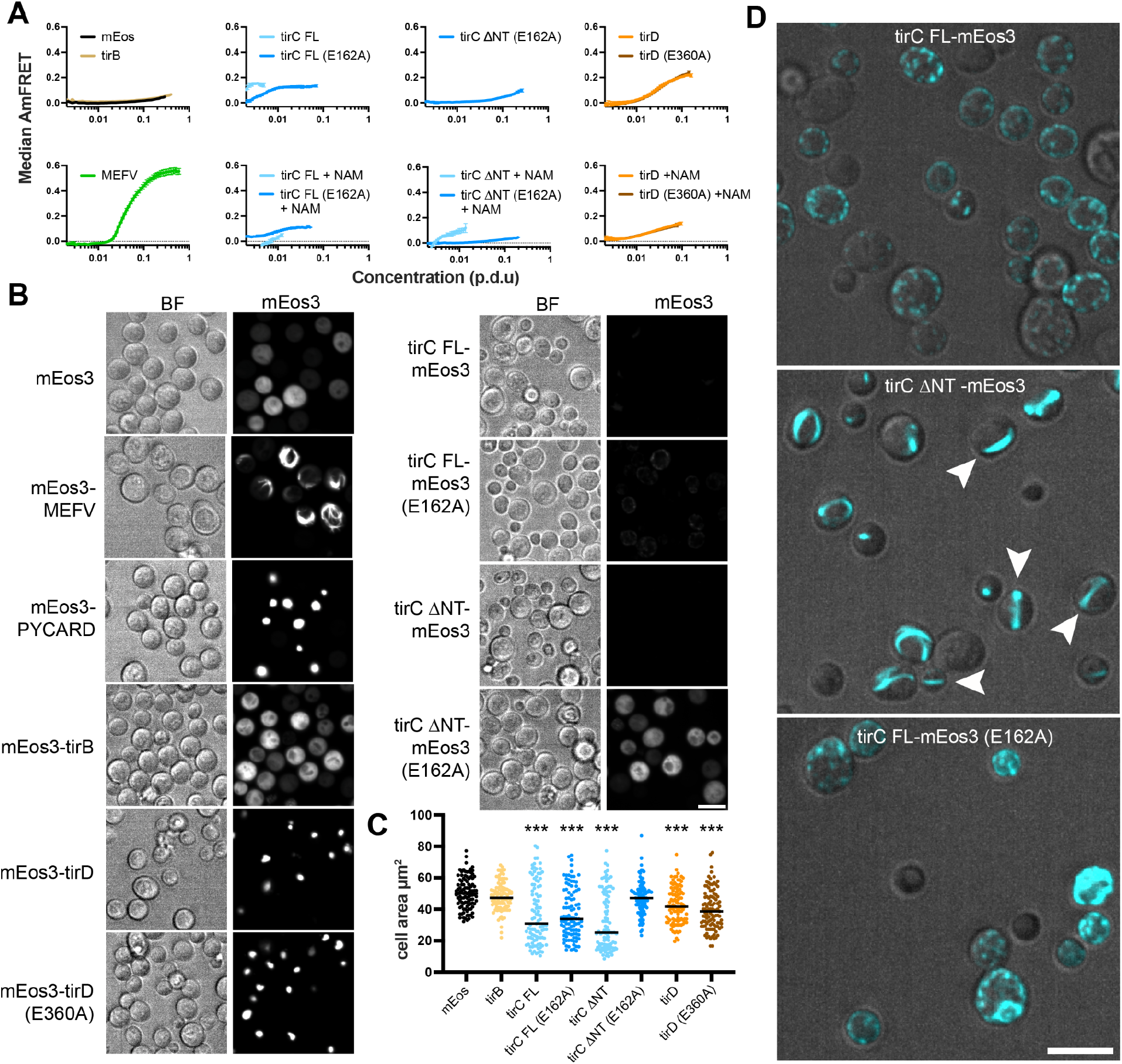
TirC and TirD form filaments and aggregates in eukaryotic cells. **A.** Splines showing median AmFRET signal from mEos3-tagged TirBCD constructs expressed in yeast cells with or without excess nicotinamide (NAM). mEos3 alone is included as a negative control, while MEFV is a positive control for filament formation. Protein concentration per cell is represented on the x-axis in procedure-defined units (p.d.u., see methods). FRET signal indicating protein self-assembly is shown as a higher signal on the y-axis. The tirC ΔNT protein did not express to detectable levels in standard conditions, but expression was partly rescued by the addition of excess NAM to the media. Error bars show standard deviations. **B.** Microscopy of yeast cells expressing mEos3-tagged proteins, with channels showing brightfield images (BF) and fluorescent signal (mEos3). Exposure and contrast levels were matched across samples. Both mEos3-MEFV and mEos3-PYCARD are positive controls for aggregation of eukaryotic immune proteins. **C.** Cell size quantification of yeast expressing TIR constructs. Black lines show median values. *** = p<0.001 when compared to the mEos control by one-way ANOVA and with Dunnett’s posttest. **D**. Microscopy images of mEos3 tagged tirC, shown with enhanced contrast to visualize low protein levels. Cells were exposed to excess NAM to partly rescue TirC expression. Arrowheads show locations of linear filaments in the TirC ΔNT sample that appear to cross the cell and/or wrap around the cell periphery. The TirC full-length (FL) proteins localized to discrete puncta in both the WT and E162A proteins. Scale bars are 10 µm.

**Figure 5.**
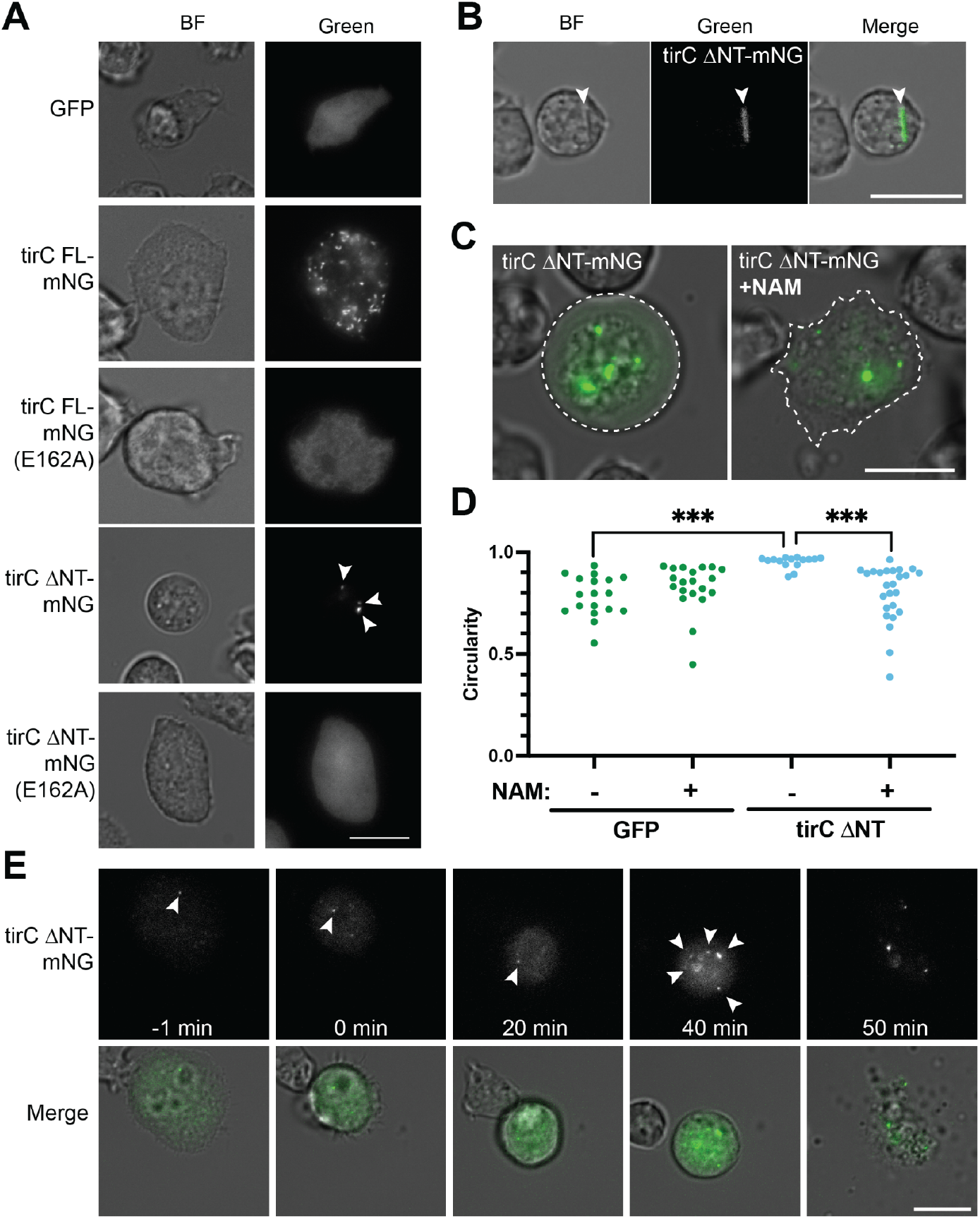
*Dictyostelium* amoebae tolerate full length TirC oligomerization, but truncated TirC is toxic. **A.** Microscopy of amoeba cells expressing GFP controls or TirC proteins tagged with mNeonGreen (mNG), showing brightfield (BF) and fluorescent (Green) channels. GFP cell lines are controls showing diffuse cytoplasmic localization. TirC proteins were either full length (FL) or truncated to lack the N-terminal helix (ΔNT). All images in A show signal from inducible constructs with matched exposure and contrast levels. Other panels have enhanced contrast of the green channel to visualize faint signals. **B.** TirC ΔNT-mNG sometimes localized to filaments that spanned the cells, in addition to the puncta shown in panels A, C, and F. **C.** Merged image showing localization of TirC ΔNT-mNG within amoebae, which were rounded and lost substrate adhesion. When cultured with 20mM nicotinamide (NAM) during induction, cell morphology and adhesion were partly rescued. Dotted lines show cell body morphology. **D.** Quantification of cellular circularity for amoebae expressing GFP or tirC ΔNT, either in the presence or absence of 20mM NAM. *** = p<0.001 by a Mann-Whitney test. **E.** Time lapse microscopy showing cell and protein dynamics during TirC ΔNT-mNG expression. An image taken 1 min before the start of the movie shows an adhered cell, which then rounded and lost substrate adhesion. At 40 min, the TirC ΔNT fluorescent signal formed multiple new puncta before cell lysis at 50 min. Arrowheads show the location of puncta or filaments. Scale bars are 10 μm.

To date the eukaryotic origins of the TirBCD family, we looked for TIR domain proteins in 81 species across Amoebozoa ^47,48^. We found that *tirC* originated recently, at the base of the Dictyostelia clade of cellular slime mold amoeba, and subsequently duplicated at least four times (Supp. Fig. 6). One of these duplications created *tirD* at the base of the *Dictyostelium* genus, and a second created *tirB* only in the *D. discoideum* clade. The two remaining duplications created lineage-specific *tirC* paralogs in other amoeba species. We also observed multiple losses: one for *tirB* and up to five losses for *tirC.* These patterns of *tirBCD* repeated gene duplication, divergence, and loss are consistent with evolutionary patterns frequently seen for other immune proteins in eukaryotes ^49,50^.

On our TIR domain trees, one close relative of TirBCD was *Sf*TIR-STING, a previously studied protein involved in bacterial anti-phage immunity (Fig. 1D) ^7,16^. Within infected bacteria, the STING domains of *Sf*TIR-STING bind to cyclic-di-GMP, leading to oligomerization, filament formation, and activation of the TIR domains as NADases. Pairwise comparisons of the TIR domains of *Sf*TIR-STING and *Dictyostelium* TirC showed that they are only 21.7% identical at the amino acid level. However, Alphafold predictions indicated that TirBCD proteins all possessed a similar TIR domain, including a distinctive beta-hairpin extension in the TIR’s C-C loop, similar to published experimental structures of *Sf*TIR-STING (Fig. 2A-B). In addition, the putative catalytic glutamate of *Sf*TIR-STING was conserved across TirBCD (Supp. Fig. 7). Because other CAP12/Pycsar TIR domains become active NADases through oligomerization, we used Alphafold to predict if TirB, C, and D could form homomeric or heteromeric multimers. The TIRs from all three proteins were predicted to dimerize with themselves and each of the other TirBCD TIR domains (Fig. 2D, Supp. Fig. 8). All had weaker dimerization ipTM scores with the other *Dictyostelium* TIRs from TirA and TirE. Beyond dimer formation, only TirC was predicted to form higher order oligomers, generating filament-like tetramers and hexamers (Fig. 2E). The TirC hexamer predictions were highly similar to the cryoEM structure of the *Sf*TIR-STING filament ^16^. Together, the *in silico* analyses of amoeba TirB, TirC, and TirD suggested potential functions as NADases and that TirC may be capable of forming higher order oligomers similar to their bacterial CAP12/Pycsar homologs.

Many TIR proteins include sensor domains that regulate TIR oligomerization and activation ^15^. But unlike previously studied proteins such as human TLR2, *Arabidopsis* RPS4, and *Sf*TIR-STING, the TirBCD proteins are small, with a C-terminal TIR domain and no other detectable domains (Fig. 2A-C). To determine if other portions of a bacterial operon could have been co-transferred with the TIR domain, we surveyed the TirBCD genomic loci. No bacterial genes or other immune-related domains were found near any of the homologs (Supp. Table 2). Therefore, we infer that the TIR domains were acquired alone and/or that other transferred domains have since been lost. To identify potential regulatory domains that arose in the amoebae, we investigated the TirBCD N-termini, which contained unstructured, poly-N linker regions (Fig. 2C, Supp. Fig. 7). Alphafold predicts that TirC has only a single, hydrophobic, N-terminal alpha helix, which we hypothesized was a transmembrane domain (Fig. 2C). In contrast, TirB and some homologs of TirD shared a small N-terminal domain that resembled a partial KH domain, a RNA-binding motif (Supp. Fig. 9). These findings suggest that domain shuffling has occurred within amoebae to link the TIR domains with variable N-termini. Overall, while TirBCD TIR domains are quite similar to *Sf*TIR-STING, we infer that their regulation is likely different because they do not have a STING domain to promote TIR activity nor do they have any other genetic remnants of an ancestral bacterial operon.

### TirC is a toxic NADase that oligomerizes into large complexes

To determine if *D. discoideum* TirB, TirC, or TirD might act as NADase enzymes, we expressed and purified His- and SUMO-tagged proteins to interrogate their biochemistry (Supp. Fig 10). *E. coli* carrying the TirC expression construct were not viable unless grown in the presence of excess nicotinamide, consistent with toxicity due to NADase activity. When we mutated a putative catalytic glutamate E162A in TirC, nicotinamide was no longer required to rescue toxicity (Supp. Fig. 7, Supp. Fig 10). We similarly observed no toxicity when expressing TirB or TirD. Both of these proteins were isolated as dimers from size exclusion columns as was the E162A TirC mutant (Supp. Fig. 10). In contrast, wild-type TirC could only be expressed at low levels and it eluted in the void fraction during size exclusion chromatography, suggesting that this TIR domain had spontaneously oligomerized into a large complex (Supp. Fig 10).

Within the void fraction, TirC was a highly active NADase, degrading NAD^+^ into nicotinamide and ADP ribose (Fig. 3). In contrast, TirB, TirD, and the E162A mutant TirC were inactive when incubated with NAD^+^ (Fig. 3). To test if protein activity could be inhibited by regulatory domains, we also tested a variety of TirBCD truncations to remove some or all of the N-terminal regions. The TIR domain alone of TirC caused toxicity, localized to the void fraction, and was an active NADase (Supp. Fig 10). However, no NADase activity was observed from any of the TirB or D truncations (Supp. Fig 11). It is possible that TirB and TirD are not NADases or that they require additional stimuli to trigger activation. We then performed the NAD^+^ co-incubations in the presence of PEG 8000 as a molecular crowding agent to test if these conditions would facilitate protein oligomerization and activation. Again, TirC was active while TirB and TirD were not. But surprisingly, while the TirC E162A mutant had no activity in the absence of PEG 8000, it behaved as an active NADase in the presence of PEG 8000 (Fig. 3, Supp. Fig 11). It therefore appears that TirC may use both the identified E162 residue and another part of the TIR domain to engage in NADase activity.

### TirC can form puncta and filaments within eukaryotic cells

Given the evidence that TirC can oligomerize and act as an NADase *in vitro*, we next investigated the propensity of TirBCD to form complexes within eukaryotic cells using Distributed Amphifluoric Fluorescence Resonance Energy Transfer (DAmFRET). In this assay, yeast cells expressed *D. discoideum* TIR proteins tagged with mEos3 fluorophores (Supp. Fig 12). When mEos3 undergoes partial photoconversion, the two fluorophore colors can engage in FRET if the fusion proteins aggregate or homo-oligomerize. Because the two colors occur at a fixed ratio regardless of cell-to-cell variation in protein expression level, the distribution of ratiometric FRET (AmFRET) across expression levels quantifies protein self-association propensity within eukaryotic cells.

We used DAmFRET to examine amoeba TIR protein oligomerization as compared to mammalian immune proteins that have been previously shown to self-assemble as puncta (PYCARD, also known as ASC) or filaments (MEFV, also known as pyrin) ^51^. The TirB protein was well-expressed in yeast and we observed little to no FRET signal as compared to the mEos negative control (Fig 4A, Supp. Fig. 13). The TirD protein self-oligomerized in both the WT and E360A constructs, particularly at high protein expression levels. In contrast, when we performed the same experiment with TirC, expression levels were very low for the full length TirC protein (TirC FL) and essentially undetectable for a version with the N-terminal helix removed (TirC ΔNT). However, we could partially rescue expression in these TirC samples by adding 100 mM nicotinamide (NAM) to the media (Supp. Fig. 13). Remarkably, for TirC FL the increased expression coincided with reduced AmFRET signal suggesting that NAM inhibits self-assembly (Fig. 4A, Supp. Fig. 13). The E162A mutant greatly increased expression for both TirC constructs, consistent with inactivation of toxic NADase activity. Self-assembly of the E162A mutants was also reduced compared to WT TirC, implying that this residue is important for oligomerization within cells, just as it was *in vitro* (Fig. 4A, Fig. 3, Supp. Fig 11).

Next, we used microscopy to observe the cellular localization of the mEos3-tagged constructs. In the samples without FRET signal (mEos3, TirB, TirCΔNT E162A), the mEos fluorescent signal appeared diffuse throughout the cells (Fig. 4B, Supp. Fig. 14). In contrast, both TirD WT and E360A proteins localized to large cellular puncta, similar to PYCARD. Because the other TirC proteins were expressed at very low levels, we altered the contrast settings and imaged these proteins with and without excess NAM to observe their localization. In the absence of NAM, we observed large filaments or aggregates in WT TirC FL-expressing yeast (Supp. Fig. 14). These became smaller and more dispersed in the presence of NAM (Fig. 4D). The full-length E162A mutant formed irregular puncta with or without NAM. Notably, the TirCΔNT protein formed puncta in the absence of NAM, and long, linear filaments that spanned the yeast cells in the presence of NAM (Fig. 4D, Supp. Fig. 14). The TirC ΔNT (E162A) samples were strikingly different, with high expression and diffuse localization similar to the mEos negative control, regardless of NAM presence (Fig. 4B). All strains carrying proteins that formed puncta or filaments (i.e. all of the TirC proteins except for TirCΔNT E162A) also showed a reduction in median yeast cell size as compared to the mEos control, reflecting toxicity at the level of cell growth (Fig. 4C).

Because the NT of TirC was predicted to be a transmembrane helix, we tested if this region impacted membrane localization and puncta formation. We expressed the TirC-mEos proteins in the presence of an orthogonal fluorescent marker that localizes to the endoplasmic reticulum (ER) and vacuole (Elo3-mTagBFP2 ^52^). Indeed, the FL TirC protein colocalized with this marker at the ER, and this was disrupted by ΔNT (Supp. Fig. 12C), suggesting that the NT targets the protein to membranes.

Finally, as an orthogonal measure of toxicity, we spotted serial dilutions of the yeast cells on media either repressing (glucose) or inducing (galactose) expression of the proteins. Similar to *E. coli,* both TirC constructs were highly toxic when expressed in yeast and this toxicity depended on TirC E162 (Supp. Fig 12B). Expression of TirD was also mildly toxic, with an approximately 3-fold reduction in cell growth. However, TirD toxicity did not depend on the homologous glutamate E360. TirB expression had no impact on yeast viability. Taken together, these data indicate that TirC can oligomerize into puncta and filaments when expressed in yeast, and both oligomers cause cell death akin to the action of *Sf*TIR-STING. Moreover, both TirC’s N-terminus and putative catalytic residue E162 promote TirC self-assembly and toxicity.

### Full length TirC is tolerated in amoebae while truncated TirC is toxic

While the toxicity, NADase activity, and oligomerization of TirC was largely consistent with the activities of its *Sf*TIR-STING homolog, we wanted to know the roles of TirBCD in their native cell context. Using CRISPR/Cas9, we generated single, double, and triple mutants in *Dictyostelium* amoebae. For each gene, we created large deletions and sometimes frameshift mutations, each of which disrupted the TIR domain (Supp. Fig. 7). When we grew each mutant in rich media, we observed no differences in growth as compared to the wild type AX2 amoebae, implying that TirBCD are not necessary for core cellular processes (Supp. Fig. 15A). Because TIR domains in other species play roles in programmed cell death, we then investigated multicellular development, a process that requires the regulated death of stalk cells to create mature fruiting bodies ^53^; ^30^. However, we did not observe any large differences in fruiting body morphology or developmental timing (Supp. Fig 16). We next asked whether TirBCD altered amoeba responses to other microorganisms as part of an immune defense. While *D. discoideum* viruses have not yet been isolated or described, the amoebae routinely sense and phagocytose bacteria. Depending on the bacterial species, these bacteria can be food for the amoebae (e.g. *K. pneumoniae*) or can survive and replicate as intracellular pathogens (e.g. *L. pneumophila*). On lawns of *K. pneumoniae* food bacteria, *tirC-* mutants created modestly smaller plaques on bacterial lawns, with *tirBC- and tirBCD-* mutants behaving similarly (Supp. Fig 15B & C). When we tested multiple, independent *tirC-* mutants in the same assay, all made smaller plaques than wild type, but only 2 out of 3 independent clones showed statistically significant reductions (Supp. Fig 15D). During *L. pneumophila* infections, we observed modestly decreased bacterial replication within *tirC-, tirBC-*, and *tirBCD-* amoeba mutants, although again there was variation across clones (Supp. Fig 15E-H). Gentamicin protection assays showed that *tirC-* mutants had reduced phagocytosis of *L. pneumophila* at the beginning of the experiment, likely accounting for the reduced intracellular replication (Supp. Fig 15I). These results hint that TirC may play a role in amoeba immunity by regulating phagocytosis, although the observed defects were mild and variable across mutant clones.

It is also possible that amoeba TIR proteins respond to unknown amoeba pathogens or environmental conditions. We hypothesized that we might recapitulate a potential for TirC oligomerization and toxicity in amoeba cells through inducible overexpression, similar to the *E. coli* and yeast experiments. We generated extrachromosomal plasmids to express mNeonGreen-tagged, full length (FL) and ΔNT TirC WT and E162A proteins in amoebae under the control of a tetracycline (Tet) responsive element. We used these plasmids to transform a strain of *D. discoideum* containing a Tet OFF transactivator. In this system, TirC expression is suppressed when doxycycline is present and protein expression is induced when doxycycline is removed. Meanwhile, another drug selects for maintenance of the extrachromosomal vector. To control for negative effects on cell growth due to overexpression and selection, we transformed the same cell line with constitutive and doxycycline-inducible GFP expression vectors in parallel. We first monitored *D. discoideum* growth and viability during TirC induction, finding similar growth rates across all of the overexpression strains (Supp Fig. 17). However, fluorescence microscopy revealed that the percentage of mNeonGreen-positive cells was low for all TirC constructs (<2%, Supp Fig. 17). Therefore, the low level of induced expression may have obscured impacts of TirC on cell viability at the population level.

Nevertheless, we could image the cells to observe cell morphology and protein localization (Fig. 5). As expected, the GFP controls showed a diffuse cytoplasmic signal. In contrast, full length TirC-mNG localized to numerous bright puncta within the cells (Fig. 5A). TheTirC FL (E162A) mutant showed more diffuse localization than TirC WT, consistent with disrupted oligomerization (Fig. 5D). Additionally, unlike the small cell sizes observed in yeast (Fig. 4C), in amoebae TirC FL overexpression did not alter cell size or morphology (Supp Fig. 17). Indeed, live timelapse microscopy of amoebae expressing FL TirC-mNG showed that expression could persist over at least 42 hours (Supp. Movie 1). During this time, the amoebae were motile, engaged in cell division, and were indistinguishable from neighboring mNeonGreen-minus amoebae, illustrating their tolerance of full length TirC expression.

In contrast, expression of TirC ΔNT induced rounding and cell death. Within amoebae, TirC ΔNT localized either in small puncta or in long filaments spanning the cells (Fig. 5A-C). These cells exhibited signs of stress such as rounding and loss of substrate attachment (Fig. 5D). Some amoebae appeared to be swelling, with cell membranes encompassing a translucent region beyond the core cell contents (Fig. 5C, Supp Fig. 17). Timelapse microscopy further revealed cell lysis events (Fig. 5E, Supp. Movie 2), which could occur immediately after rounding or several hours later. Remarkably, some of the TirC ΔNT-mNG puncta remained intact following explosive cell lysis, demonstrating the stability of these assemblies. The timelapse imaging also showed that oligomerization was dynamic, with new puncta appearing over time (Fig. 5E). In comparison, the TirC ΔNT (E162A) protein was diffusely cytoplasmic and non-toxic in all conditions, similar to the GFP controls (Fig 5A, Supp Fig. 17).

To test if the amoeba rounding and lysis was due to NAD depletion, we induced TirC ΔNT-mNG expression in the presence of nicotinamine. Indeed, this condition rescued the amoeboid cell morphology and restored attachment in TirC ΔNT-expressing cells (Fig. 5C-D, Supp. Fig 17). Taken together, these data suggest that full length TirC can oligomerize into puncta in *D. discoideum*, yet the amoeba cells somehow tolerate TirC FL expression and oligomerization, unlike heterologous systems. However, TirC ΔNT toxicity mirrored the defects seen in yeast and bacterial expression, which likely derives from NAD^+^ depletion.

Because TirC FL expression was tolerated in amoebae but not other organisms, we hypothesized that other factors in the amoeba cell might be used to prevent autoactivation of TirC’s NADase activity. Given the potential for TirB and/or TirD to dimerize with TirC (Fig. 2D), we asked if these paralogs could serve as TirC inhibitors. However, within *in vitro* experiments mixing together all combinations of TirB, TirC, and TirD purified proteins, all TirC-containing mixes behaved identically to TirC alone (Supp. Fig 18). In addition, when we performed pairwise co-expression of all the proteins within yeast cells, we did not observe any enhancement or suppression of TirC toxicity (Supp. Fig 19). Therefore, we propose that the amoebae have successfully domesticated the TIR domain of TirC to regulate its activity, but this mechanism appears to be independent of TirB and TirD.

## Discussion

Newly acquired genes can only be useful if they are safely incorporated into a cell’s existing molecular and cellular processes. Several pieces of evidence suggested that this domestication has happened for TirBCD following the horizontal gene transfer of a TIR domain from prokaryotes into eukaryotes (Fig. 1 & 2). At the genetic level, *tirBCD* genes are similar to other *D. discoideum* loci in their gene structures (e.g. intron acquisition for *tirB* and *tirD*), GC content, sequence repeats, distinct patterns of gene expression, and evolutionary history of gene duplication and divergence (Supp. Fig 5 & 7). Therefore, these proteins appear to have lost their prokaryotic (and gained eukaryotic) gene regulation. Overexpression of full length TirC is also tolerated within *D. discoideum* cells, causing minimal disruptions in cellular behavior despite protein oligomerization into discrete cellular foci (Fig. 5, Supp. Movie 1). This tolerance is in stark contrast with the strong, spontaneous NADase activity we observed from TirC’s TIR domain *in vitro,* the toxicity of TirC ΔNT in amoebae, and thetoxicity of both TirC FL and ΔNT when expressed in bacteria or yeast (Fig. 3, 4, 5, Supp. Fig 10, 12, Supp. Movie 2). In these situations, even very low concentrations of TirC were toxic: leaky expression in *E. coli* inhibited bacterial growth even in uninduced conditions, while in the yeast DAmFRET system TirC oligomerized and was toxic at the lowest detectable cellular concentrations (Fig. 4, Supp. Fig 10, 13). In all cases, TirC oligomerization was dependent on the putative catalytic residue E162. Based on these results, we infer that the acquisition of the N-terminal, transmembrane helix was essential for the amoeba domestication of TirC’s NADase activity, but the addition of this domain is not sufficient to to prevent TIR NADase autoactivation when the protein is moved into other organisms.

These findings illustrate the potential dangers to eukaryotes of acquiring antiphage proteins through horizontal transfer and altering their mechanisms of regulation. While this process has the potential to give rise to new aspects of eukaryotic cell biology and innate immunity, new proteins can come with a high potential for toxic side effects. Even within bacteria that have intact defense operons, immune defenses are costly, leading to high rates of gene loss ^12^. The costs are likely even higher when regulation is disrupted during HGT ^11^. Previously, these aspects have been difficult to measure because many instances of bacteria-eukaryote HGT are extraordinarily ancient, sometimes dating back over one billion years ^6,54,55^. This vast time scale can make it difficult to reconstruct the steps that forged the links between bacterial and eukaryotic immunity. But because of its unique timeline, the TirBCD system allows for a more direct interrogation of the process of immune gene domestication.

One prediction of shared immune defenses is that the catalytic activity of eukaryotic homologs will be much lower than that of the prokaryotic ancestors ^11^. We note that TirC’s enzymatic activity is strong and its toxicity has notable parallels to bacterial TIR proteins that deplete cellular NAD^+^ ^7,56^. During TirC overexpression, it also readily oligomerizes into large complexes in bacterial lysates, in amoeba cells, and in yeast cells, where we observed long, linear filaments reminiscent of *Sf*TIR-STING (Fig. 4, 5, & Supp. Fig 10). Thus, it appears that much of the biochemical activity of the TIR domain has been retained following transfer, despite the differences in regulation when TirC is moved across taxa.

In future studies, it will be very interesting to discover in detail how amoebae avoid the spontaneous toxicity of TirC. Although TirB and TirD were logical candidates, we found no evidence that these proteins altered TirC NADase activity or toxicity (Supp. Fig. 17 & 18). Instead, we speculate that amoebae have used TirC to co-opt a form of bacterial cell death for use in eukaryotes, wherein a cytoplasmic TIR domain oligomerizes and depletes cellular NAD^+^. While potential triggers remain unknown, it is easy to imagine that proteolytic cleavage (mediated by either amoeba or pathogen proteases) could release the TIR domain from its transmembrane tether, leading to rapid activation. If so, TirC would mirror important aspects of mammalian inflammasome activation via protease activity in NLRP1 ^57^. Alternatively, TirC could be activated if cells produced an alternate form of the transcript, lacking the N-terminal helix. Amoebae may also use other endogenous proteins to bind TirC and prevent TIR domain activation within TirC FL puncta (Fig. 5A). Beyond TirC, we expect that TirB and TirD are important for amoeba TIR signaling but may involve novel mechanisms, given that TirD oligomerized and was toxic in yeast but both activities were independent of the putative catalytic glutamate. Ultimately, these future studies will reveal how organisms avoid the consequences of acquiring potentially toxic modules, illustrating overall how innovations in immunity arise.

This detailed study investigating how eukaryotes modify and deploy horizontally acquired immune genes was made possible because of the expansive phylogenetic analyses of TIR domains (Fig. 1). These studies revealed the close relationships between prokaryotic TIRs and amoeba TirBCD. While some aspects of TIR diversity have been characterized across bacterial species ^34,58^, a global view of eukaryotic TIRs has been lacking. The new categorization scheme for TIRs presented here can help to identify and characterize members of this large, diverse protein family. Some identified clades such as clades 10-13 are widespread in eukaryotes, but unstudied because they are found outside of traditional model organisms. Based on precedents from bacterial, plant, and animal TIRs, we hypothesize that these highly diverged eukaryotic proteins are likely to oligomerize and exhibit a wide array of novel biochemical functions in their regulation of immunity and/or cell death. We hope that the TIR Atlas and phylogenetic tools will be of use to the field as we continue to explore this expansive protein diversity.

## Author contributions

**EMC**: Conceptualization, Data Curation, Formal analysis, Investigation, Methodology, Resources, Supervision, Validation, Visualization, Writing- original draft, Writing- review & editing; **ECL**: Conceptualization, Data Curation, Formal analysis, Investigation, Methodology, Resources, Supervision, Validation, Visualization, Writing- original draft, Writing- review & editing; **JLP**: Conceptualization, Data Curation, Formal analysis, Investigation, Methodology, Resources, Validation, Visualization, Writing- original draft; Writing- review & editing; **MH**: Data Curation, Formal analysis, Investigation, Resources, Validation, Visualization, Writing- original draft, Writing- review & editing; **JRD**: Data Curation, Formal analysis, Investigation, Resources, Validation, Visualization, Writing- original draft; **JJL**: Investigation, Methodology, Visualization; **YL**: Investigation, Methodology; **NG**: Investigation; **HG**: Investigation; **BRM**: Conceptualization, Investigation, Methodology, Resources, Supervision, Visualization, Writing- review & editing; **RH**: Conceptualization, Investigation, Project administration, Resources, Supervision, Visualization, Writing- original draft; Writing- review & editing; **TCL**: Conceptualization, Funding acquisition, Investigation, Methodology, Project administration, Resources, Supervision, Visualization, Writing- original draft; Writing- review & editing

## Materials and Methods

### Strains, Plasmids, and Primers

Detailed information about all strains (*E. coli, S. cerevisiae, D. discoideum, K. pneumonia,* and *L. pneumophila*) is located in Supplementary Table 3. Plasmid information is located in Supplementary Table 4. Primers used in this study are found in Supplementary Table 5.

### Iterative HMM searches and construction of phylogenetic trees

#### 1. Eukaryotic TIR search strategy

We searched for TIR domain sequences in Eukprot v3 similar to previous protocols ^6^. We started each search from the published PFAM HMMs from the STIR clan, beginning with PF01582 and then searching in this order: PF13676, PF18567, PF08937, PF10137, PF08357. We continued iterating the searches while broadening the HMMs until one of these criteria were met: 1) a minimum of two searches per PFAM HMM were performed, 2) a search failed to find unique TIR domains, or 3) all new proteins found were unlikely to be TIR domains as determined by qualitative visual comparison between *bona fide* TIR domains and structural predictions made via Alphafold (see AlphaFold model predictions and TIR domain assessment). This resulted in between 2-5 eukaryotic searches for all five Pfam starting groups. With the newly expanded HMMs, we then used hmmscan (included within hmmer v3.2.1) with settings “—domtblout—domE 1e-3” to define the boundaries for each TIR domain. All eukaryotic sequences were trimmed down to their TIR domain. In cases where multiple TIR-calls overlapped, the longest sequence was kept.

We recovered 7,432 eukaryotic TIR sequences. To reduce the number used for analyses while retaining phylogenetic diversity, we next downsampled all eukaryotes except Metazoans and Amoebozoans down to 1000 sequences. To do so, the TIR domains were then aligned with MAFFT (default settings) and trimmed with TrimAL (-gt 0.1), then made into a phylogenetic tree with FastTree (default settings). Phylogenetic diversity analyzer (PDA) was then run (-k 1000). The pre-trimmed, downsampled sequences were then combined with the Metazoan and Amoeba sequences.

In our experience, EukProt tends to have a low diversity of fungal sequences and indeed we found almost zero fungal TIRs. The ones we did find were from a single species, *Rhizoclosmatium globosum* (Supp. Table 1). Within InterPro on 2025-11-02, the entire STIR clan similarly contained 0 fungal sequences. Although a very small number of fungal TIRs have previously been reported ^13^, these sequences were not included in our analyses.

#### 2. Gathering Bacterial Sequences

Bacterial sequences were gathered from Interpro from each of the five TIR Pfam families (PF01582, PF13676, PF18567, PF08937, PF10137). Each of these was trimmed down to its TIR domain, aligned with MAFFT (v7.525, default settings) and trimmed with TrimAL (-gt 0.05), then made into a phylogenetic tree with FastTree (default settings). Each of these trees was then reduced down to 100 sequences maximum. For TIR_2 (PF013676), we reduced this family of bacterial sequences using PDA down to 1000 sequences due to the vast diversity (∼29,000 sequences) in the InterPro database. Archaea from each of these groups was also extracted, aligned, trimmed, plotted on a phylogenetic tree, and reduced down to 100 sequences.

#### 3. Alignment and generation of phylogenetic trees

With the final list of 4664 TIR and SEFIR sequences we then aligned the TIR domains with MAFFT or MUSCLE (v5.1, default settings), trimmed with TrimAL (either -gt 0.10 or -gt 0.20), and constructed the phylogenetic trees with IQTree (v3.0.1, -m MFP, -bb 1000). We defined supported clades by determining which groups of sequences were found in supported (>70 bootstraps, with at least 100 sequences) clades in 3 out of 4 of the phylogenetic trees (MAFFT, MUSCLE, trim 10%, trim 20%). To focus on the largest TIR groupings, we only reported clades that retained greater than 20 sequences after removing sequences that didn’t meet the 3/4 trees support threshold.

#### 4. Alignment and generation of phylogenetic trees within individual clades

Smaller phylogenetic trees were generated via aligning the pre-trimmed TIR domains of the sequences in the smaller clades (Ex: Clade 1, Clade 4, Clade 6, and Clade 8). These were aligned with MUSCLE, trimmed with TrimAL (-gt 0.05), and constructed a phylogenetic tree with IQTree (v3.0.1, -m MFP, -bb 1000).

### AlphaFold model predictions and TIR domain assessment

AlphaFold v3 ^59^ was used extensively to generate all structural predictions. Structural predictions were used to predict potential interactions between multimeric states and with other macromolecules (Supp. Fig. 8-9). AlphaFold predicted structures were also used as an orthogonal way to support our call of which sequences were TIR domains and which were not. TIR domains have a well documented and conserved fold of five alternating alpha helices and beta sheets, and we considered models that had at least 4 out of five alpha helices and 3 out of five beta sheets as TIR domains.

### Validating TirBCD gene models in *D. discoideum*

#### 1. Discrepancy in tirC and tirD gene models across D. discoideum genomes

The gene models for *tirC* and *tirD* differed significantly in the AX2 ^43^ and AX4 ^42^ genomes. For *tirC*, the AX4 annotation predicted an early intron after the first two amino acids while the AX2 annotation predicted no introns. When we looked at the alignments of the *tirC* locus across sister species, there was extensive amino acid conservation in the predicted intronic region. We therefore hypothesized that this intron was likely called inaccurately. For *tirD,* the AX4 annotation predicted two genes (*DDB_G028737* and *DDB_G0287321*) with the first containing an intron. The AX2 annotation predicted the locus as a single gene product with an intron in the same position. The alignments of the *tirD* locus across species showed no strong support for either annotation model (AX2 or AX4) as there was no significant drop in conservation at either predicted intron, although the supposed “intergenic region” called in AX4 also showed conservation at the amino acid level. To resolve these ambiguities, we used RNA-seq and PCR of cDNA to determine the gene models for *tirB*, *tirC*, and *tirD*.

#### 2. Culturing and harvesting Dictyostelium cells

AX2 Dictyostelium discoideum were harvested for RNAseq analysis from either a bacterial lawn or axenic cultures grown in HL5 media (Axenic medium, Formedium #HLB0103, supplemented with 55.5 mM Glucose, pH 6.65) shaking at 220 rpm, 22°C. Axenic AX2 cultures were grown to a density of ∼5e^6^ cells/mL in HL5 media. Cells were pelleted and frozen at −80°C. For bacterial lawns, we grew *Klebsiella pneumoniae* overnight cultures in LB media (1% Tryptone, 0.5% Yeast Extract, 171.12 mM NaCl) at room temperature. *K. pneumoniae* cultures were then washed twice with sterile SorC buffer (14.7 mM KH_2_PO_4_, 2.04 mM Na_2_HPO_4,_ 50 µM CaCl_2_, pH 6.0) before their density was adjusted to an OD_600_ = 1 in SorC. Axenic AX2 cells were washed twice in SorC before being counted on a hemocytometer and their density adjusted to 1e^6^ cells/mL. We added 100 μL of AX2 cells to 600 μL of *K. pneumoniae* and plated 600 μL of this solution onto SM/5 plates (11.1 mM glucose,0.2% proteose peptone, 0.02% yeast extract, 0.83 mM MgSO_4_, 2.79 mM KH_2_PO_4_, 0.688 mM K_2_HPO_4_, 2% agar, pH 6.2). We incubated the plates for 3 days at 22°C before harvesting the amoeba cells off the plates with a cell scraper. AX2 cells were enriched by centrifugation of harvested cells at 100 x g for 1 minute through 5 mL of 20% Percoll (Fisher Scientific 45-001-748) diluted in PBS to separate the lighter *K. pneumoniae* cells from the heavier *D. discoideum* cells. We repeated this percoll separation three times per sample. *D. discoideum* cells were then frozen at −80°C.

#### 3. RNA extraction, DNase treatment, and RNAseq analysis

RNA was extracted from *D. discoideum* cell pellets via RNeasy RNA extraction kit (Qiagen 74104). Purified RNA was then treated with TURBO DNase (ThermoFisher Scientific AM2238). Purified, DNA-free, RNA was then sequenced via illumina sequencing (Plasmidsaurus Inc.). Data quality was assessed using FastQC (v0.12.1). Reads were then quality filtered using fastp (v0.24.0) with these parameters: poly-X tail trimming, 3’ quality-based tail trimming, a minimum Phred quality score of 15, and a minimum length requirement of 50 bp. The remaining reads were aligned with STAR aligner (v2.7.11) to the AX2 genome followed by coordinate sorting using samtools (v1.22.1). UMIcollapse (v1.1.0) was then used to remove PCR and optical duplicates. Filtered and de-duplicated reads from three biological replicates were mapped onto the AX2 genome in Geneious Prime for visualization. Coverage was calculated as the number of reads that mapped at each base pair on the sequence.

#### 4. Genomic DNA extraction, cDNA synthesis, and PCR

Crude genomic DNA was isolated from AX2 *Dictyostelium discoideum* that were grown in axenic culture in HL5 media (22°C). Cells were pelleted and resuspended in lysis buffer ^60^ (50 mM KCl, 10 mM Tris pH 8.3, 2.5 mM MgCl_2_, 0.45% IGEPAL CA-630, 0.45% Tween 20) with 0.6 mg/mL Proteinase K (ThermoFisher Scientific FERE0491) and incubated at 60°C for 1 hour and then 95°C for 15 minutes.

#### 5. Inferred gene models

Because the *tirB* gene model was identical across AX2 and AX4 genomes, this gene served primarily as a positive control in our RNAseq mapping and cDNA PCR experiments. The RNA-seq reads spanned the exon-exon junction at the exact expected coordinates. Our cDNA PCRs for *tirB* also showed the expected bandshift following intron splicing, confirming the presence and size of the intron, as well as validating that the cDNA was free of genomic DNA. In the RNA-seq reads, a small minority of reads mapped to the predicted intron-exon boundary of *tirB* suggesting that a small amount of genomic DNA contamination may have been present (Supp. Fig. 4A). Overall, the data supports the existing *tirB* gene model.

The *tirC* locus was not strongly expressed in our samples. Of the reads that mapped, we observed RNAseq reads that mapped to the putative AX4 intronic region but did not observe any intron spanning reads. However, the cDNA PCRs for *tirC* showed a single band that was the same size as the genomic DNA amplified band, supporting the AX2 gene model annotation that encodes *tirC* as a single exon (Supp. Fig. 4B).

The *tirD* RNA-seq coverage was even lower. There was no coverage over a large section of the gene that is predicted to be an exon in both gene models. At the 5’ end, most of the reads supported the *tirD* intron that was predicted by both the AX2 and AX4 annotations. However, a minority of reads mapped to the predicted exon-intron boundaries or that spanned the intron with a different splice donor. The cDNA PCR that used primers crossing the two predicted AX4 genes generated a single mRNA product, supporting that they are part of the same transcript (Supp. Fig. 4C). Thus, *tirD* appears to match the AX2 gene annotation, with the possibility of additional splice isoforms.

### Search for bacteria-derived genes near *tirB, tirC, tirD* loci

The neighboring genes (3 upstream, 3 downstream) of the *tirB, tirC,* and *tirD* locus of *Dictyostelium discoideum* strain AX2 were queried with nBLAST searches against the Core nucleotide database (core nt) excluding results from Dictyostelia (taxid: 33083). In supplemental table 2 we report the top hit for each gene (E-value < 0.05).

We next took the protein translations of these neighboring genes and first used hmmscan (hmmer v3.2.1) with settings “—domtblout—domE 1e-3” against the Pfam database (Pfam-A.hmm) to search for the domains in each protein. We then searched for homology to other proteins with Jackhmmer (accessed https://www.ebi.ac.uk/Tools/hmmer/search/jackhmmer on 4/21/2026) using default settings, searching against the Reference Proteomes (2025_01). We recorded the taxonomic classification of the significant hits (E-value cutoff 0.01) at the domain level.

### TirBCD history across amoeba species

To search for the evolutionary history of TirBCD across amoeba species, we took a closer look at all the available amoeba genomes that we could search for TirBCD-like homologs. The Eukprot database includes genomes or transcriptomes from 72 amoeba species within Amoebozoa, including 13 from the Eumycetozoa clade containing *Dictyostelium discoideum*. We expanded our search to the genomes of 10 recently published *D. discoideum* isolates and sister species ^43^ as well as *Polysphondylium violaceum* (Genbank: AJWJ01001228.1), as they are not represented in Eukprot v3. To identify TirBCD homologs, we searched with every TIR HMM that we had generated during our iterative HMM searches and then used AlphaFold to validate whether or not these sequences folded like TIR domains. To identify the origin of TirBCDs, we therefore looked in more closely related amoeba species using tblastn. We searched custom databases containing the genomes from fifteen Dictyostelia isolates that branched outside of *D. intermedium* (Supp. Fig 6) using tblastn with a cutoff of 1e^-5^. Across all Dictyostilia TirBCD homologs, we made alignments using MAFFT and constructed phylogenetic trees using PhyML to categorize sequences as *tirB, tirC,* or *tirD*.

### TirBCD expression in *E. coli* and protein purification

All the genes were cloned using Gibson assembly into linearized vectors and transformed into Top10 *E.coli* cells, proper gene insertion was confirmed with Sanger sequencing, and the plasmids were then transformed into BL21(DE3)-RIL *E.coli* cells for protein expression. Colonies from TirB, TirC, TirC (E162A), and TirD were grown on MDG-agar solid media (0.5x trace metal mix [60 mM HCl, 50 mM FeCl_3_*6H_2_O, 20 mM CaCl_2_*2H_2_O, 10 mM MnCl_2_*4H_2_O, 10 mM ZnSO_4_*7H_2_O, 2 mM CoCl_2_*6H_2_O, 7H_2_O, 2 mM CuCl_2_*2H_2_O, 2 mM NiCl_2_*6H_2_O, 2 mM Na_2_MoO_4_*2H_2_O, 2 mM Na_2_SeO_3_, 2 mM H_3_BO_3_], 2 mM MgSO_4_, 0.5% glucose, 1x M solution (1.25 M Na_2_HPO_4_, 1.25 M KH_2_PO_4_, 2.5 M NH_4_Cl, 0.25 M Na_2_SO_4_), 0.25% Aspartic acid, 100 μg/μl ampicillin, 34 μg/μl chloroamphenicol). TirC was grown on and in MDG media supplemented with 20 mM nicotinamide (NAM) to mitigate toxicity. Selected single colonies were grown overnight at 37°C in 30mL MDG liquid media for 16-20 hours and shaking at 230 RPM. The overnight culture was used to inoculate 1L of M9ZB media (47.8 mM Na_2_HPO_4_*7H_2_O, 22 mM KH_2_PO_4_, 18.7 mM NH_4_Cl, 85.6 mM NaCl, 1% cas-amino acids, 0.5% glycerol, 2 mM MgSO_4_, 0.5x trace metal mix, 100 μg/μl ampicillin, 34 μg/μl chloramphenicol) in 2.5L flasks. After the cultures reached an OD_600_ greater than 2.0 the flasks were placed on ice for 20 minutes, 0.5 mM of IPTG (final) was added to the flasks and incubated at 16°C shaking at 230 RPM overnight. The cells were harvested by centrifugation, washed with chilled PBS buffer and flash frozen with liquid N_2_ and stored at −80°C until needed.

Nickel affinity chromatography using gravity flow at 4°C was carried out as a first step purification. The *E.coli* pellets were resuspended in lysis buffer (20 mM HEPES pH 7.5, 400 mM NaCl, 10% glycerol, 30 mM imidazole, and 1 mM DTT) and sonicated (10 seconds on, 20 seconds off at 70% amplitude for 5 minutes total on-time). The cell debris was subjected to centrifugation, and the clarified lysate was applied to 7 ml of packed Ni-NTA resin equilibrated with lysis buffer. The column was washed with 20 mL of lysis buffer, 70 mL of wash buffer (lysis buffer supplemented to 1M NaCl) and 35 mL of lysis buffer. The protein was eluted with 20mL of elution buffer (lysis buffer supplemented to 300 mM imidazole). The elution fraction was dialyzed overnight in dialysis buffer (20 mM HEPES pH 7.5, 250 mM KCl, 1 mM DTT, and 5-10% glycerol) at 4°C with hSENP2 SUMO protease to cleave the SUMO2 solubility-tag (if present). The dialyzed elution was spin-filter concentrated and loaded onto a Hiload Superdex 75 or HiLoad 16/600 Superdex 200 pg (Cytiva) size exclusion column equilibrated with gel filtration buffer (20 mM HEPES pH 7.5, 250 mM KCl, 1 mM TCEP-KOH). Fractions that contained the protein of interest were collected and analyzed with SDS-PAGE. Fractions were pooled and concentrated to >10 mg/mL, proteins <10 mg/mL were stored with 10% glycerol. The proteins were flash frozen in liquid N_2_ and stored at −80°C.

### HPLC analyses

Enzymatic reactions were performed in a total volume of 150 μL with 500 μM NAD^+^, 100 mM NaCl, 20 mM HEPES-KOH (pH 7.4), +/- PEG 8000 (10%), with 20 μM of protein added last. The concentration for TirC was estimated because we could not get consistent purity by SDS-PAGE. Reactions were incubated at room temperature overnight (16-20 hrs) and then transferred to 10 KDa cut-off spin-filter and centrifuged for 20 min at 13,500g. 10μl of all samples were injected onto an Agilent 1200 series equipped with a 4.6 × 150 mm and 5 μm particle-size Zorbax Bonus-RP column using an isocratic elution method including 97% 50 mM NaH_2_PO_4_ (pH 6.8) and 3% acetonitrile buffer system held at 40°C. The reaction components were monitored at 254 nm using an in-line UV absorbance detector.

Time course enzymatic reactions for TirC and TirC_E162A were performed in a total volume of 550 μL with 500 μM NAD^+^, 100 mM NaCl, 20 mM HEPES-KOH (pH 7.4), +/- PEG 8000 (10%), with 20 μM of protein added last. Reactions were incubated for 0-3 hours, after which 100 μL of the reaction mixture was transferred to a new tube and heat-inactivated for 5 minutes at 95 °C. The sample was then transferred to 10 KDa cut-off spin-filters and centrifuged for 20 min at 13,500g. Filtered reactions were then analyzed with HPLC as described above.

TirB/C/D mixing experiments were performed in a total volume of 100 μL with 500 μM NAD^+^, 100 mM NaCl, 20 mM HEPES-KOH (pH 7.4), with 20 μM of each protein added last (TirC concentration estimated as with all other reactions). Reactions were incubated for 2 hours then heat inactivated for 5 minutes at 95 °C. The samples were then transferred to 10 KDa cut-off spin-filters and centrifuged for 20 min at 13,500g. Filtered reactions were then analyzed with HPLC as described above.

### TirBCD expression in yeast and DAmFRET

Yeast strains were prepared as described in Venkatesan et al. (2019) ^61^ and Khan et al. (2018) ^62^. DAmFRET data collection and analysis were as previously described ^63^ except with the addition of 10 or 100 mM nicotinamide where indicated during induction. Briefly, proteins of interest were yeast codon-optimized and cloned behind the *GAL1* promoter in high-copy vectors V08 and V12 for C- and N-terminal tagging, respectively, with mEos3.1. Single parental yeast strains for each protein of interest were made via a standard lithium-acetate transformation protocol using yeast strain rhy2054. Individual colonies were picked in biological triplicate and incubated in 200 μl of a standard synthetic media containing 2% dextrose (SD-ura) overnight while shaking on a Heidolph Titramax-1000 at 1000 rpm at 30°C. Following overnight growth, cells were resuspended in media containing 2% galactose (SGal-ura) to induce ectopic protein expression and treated with either 0, 10 mM, or 100 mM nicotinamide (Sigma, N0636) and then returned to shaking incubation for 16 h. Cells were resuspended in fresh inducing media 4 h before photoconversion to reduce autofluorescence. For DAmFRET analysis, microplates containing yeast cells were photoconverted using OmniCure S2000 Elite fitted with a 320–500 nm (violet) filter and a beam collimator (Exfo), positioned 45 cm above the plate, to deliver an irradiance of ∼12 J/cm^2^ over 3.72 min while shaking at 1000 rpm. Immediately following photoconversion, yeast cells were assayed on a flow cytometer (BioRad ZE5 Analyzer).

The ratio of Acceptor expression (Acceptor-A) to Side scatter (SSC-A) served as a proxy for protein concentration^64^. To visualize the relationship between AmFRET and protein concentration, data were partitioned into 64 concentration bins (p.d.u.). A "spline" was constructed by smoothing the median AmFRET values across these bins as described previously ^65^. Low-concentration bins were excluded where signals fell below the assay’s limit of detection. Final plots represent means +/- SD of triplicates.

### Fluorescence microscopy of yeast cells

Replicate transformant colonies of yeast were prepared for confocal imaging using the DAmFRET protocol as described above, with the exception of photoconversion. Cells were transferred to a PhenoPlate-96 optical microplate (Revvity, 6055302) for imaging on the Opera Phenix high-content screening system (Revvity, formerly PerkinElmer). Confocal images were acquired using a 40x water objective lens (N.A. 1.1). The green form of mEos3.1 was excited using a 488 nm laser and emission was collected with 500-550nm bandwidth emission filters.Transmitted light was also collected. Z stacks were acquired for a total distance of 0.8 μm, with a spacing of 0.2 μm and each channel was max projected (NumPy) in the Z plane to obtain the final images. Four fields of view were acquired for each sample. For image analysis, images were processed using Fiji (ImageJ, v1.54p). Brightness and contrast were adjusted based on the expected brightest biological sample, and those settings were propagated to all other images within the panel, as in Fig. 4. Due to the large variation in mEos3.1 signal intensity between samples, brightness and contrast were visualized at 2 levels in Supp. Fig. 14. For size quantification of yeast cells, a random sampling of cells was obtained using two separate transmitted light images for each sample for a total of approximately 100 cells per sample. Regions of interest (ROIs) were hand drawn around the perimeter of each cell using the ellipse tool in Fiji. These ROIs were then used to measure the area of each cell based on the set scale.

### Yeast toxicity assay

*URA3*-marked plasmids encoding the proteins of interest tagged with mEos3.1 were transformed into parental yeast strain rhy3332. A separate set of *LEU2*-marked plasmids encoding the same proteins of interest tagged with BDFP1.6:1.6 were transformed into parental yeast strain rhy3333. Both strains contained a *tTA{Off}tetO7^WHI5_hphMX* construct replacing endogenous *CLN3* such that doxycycline (dox; 40 µg/ml) is required for cell growth. This genetic feature was incorporated to allow cell cycle arrest independently of exogenous gene expression, such that the same strains could be analyzed by both serial dilutions and DAmFRET as desired (though not utilized for DAmFRET in this study). The transformed strains were then mated in pairwise combination and selected on SD-ura-leu+dox to obtain diploid cells harboring both plasmids. Cells were inoculated into liquid SD-ura-leu+dox and grown overnight at 30 °C. Each strain was normalized to an OD_600_ of ∼1 and serially diluted 5-fold in water. Equal volumes of each dilution were immediately transferred to both SD-ura-leu+dox media plates and SGal-ura-leu+dox media plates. Plates were imaged post 72 h of growth at 30 °C.

### Generating TIR deletion mutants in *Dictyostelium* amoebae

We used CRISPR/Cas9 ^66^ to delete large regions of the *tirB, tirC*, and *tirD* loci, aiming to delete part or all of each gene’s TIR domain in single, double, and triple mutant amoeba lines. We used the Cas-Designer tool (http://www.rgenome.net/cas-designer/) to select two 20 nucleotide target sequences flanking the TIR domains during *D. discoideum* genome editing. The guide sequences and their complementary sequences were synthesized as oligonucleotides with the overhangs to allow for integration onto the CRISPR vectors. The complementary oligos were annealed to generate double stranded structures and phosphorylated. Then, a Golden Gate reaction mix was prepared with the phosphorylated guides and the CRISPR vectors pTM1544 (to deliver 2 guides per vector) or pTM1285 (to deliver 1 guide per vector). The Golden Gate product was then transformed into 5 alpha competent cells (New England Biolabs C2987). After plasmid sequencing to confirm correct assembly, the vectors with integrated guides were used to transform AX2 wild type *D. discoideum* cells via electroporation. AX2 cells were chilled then washed twice in cold, sterile EP buffer (10 mM NaPO_4_, 50 mM sucrose, pH 6.1). Cell density was adjusted to 1.38×10^7^ cells/mL in cold EP buffer. Fifteen µg of DNA was introduced into 1.0×10^7^ total AX2 cells in 0.4 cm gap cuvettes with a BioRad Gene Pulser Xcell system with the following parameters: exponential decay, 1000V, 50 ohm resistance, 50 µF capacitance. Cells were electroporated in 2 pulses with a 5 second interval. Electroporated cells were immediately allowed to recover in HL5 rich media supplemented with 50 U/mLPenicillin/Streptomycin at 22°C for ∼24 hours. After recovery, the media was replenished with HL5 supplemented with G418 [10ug/mL] and cells were incubated at 22°C for 3 days. Transformants were washed with sterile PBS to remove debris and traces of G418 and media was replenished with HL5 supplemented with Penicillin/Streptomycin.

Transformants were monitored for viability and growth over ∼7 days. Once confluent, polyclonal transformant populations were sampled to generate crude cell lysates to serve as PCR templates as described in *Genomic DNA extraction, cDNA synthesis, and PCR* methods. To genotype the transformants, we used PCR to amplify a large locus across the targeted region.

Polyclonal populations containing deletion mutants were identified as samples that contained two bands, one corresponding to the size of the intact WT locus and another significantly smaller band corresponding to a large deletion. Populations containing potential deletion mutants were isolated into monoclonal populations by limiting dilutions in HL5. Once confluent, monoclonal populations were sampled via PCR to confirm genomic editing. The amplicons were column purified (Zymo Research Corporation D4013) and sent off for Oxford Nanopore sequencing (Plasmidsaurus Inc.) to identify the deleted region. Finally, the confirmed deletion mutant cultures were expanded to make cell stocks and used for functional assays.

### Amoeba growth in rich media or in the presence of food bacteria

To test the viability of mutant and overexpression amoebae, we performed growth assays in sterile media. Amoeba were grown to confluence in t25 flasks at 22°C without shaking. Amoebae were harvested from the flasks, washed twice in SorC media and counted with a hemocytometer. Three mL cultures of HL5 in test tubes were inoculated with 5e^4^ cells/mL and grown at 220 rpm shaking at 22°C. Cells were counted on a hemocytometer at day 0 and every 24 hours for 6 days.

To determine if mutant amoebae had growth defects in the presence of food bacteria, we co-cultured these species on agar plates and looked for the formation of amoeba plaques on bacterial lawns. Overnight cultures of GFP *Klebsiella pneumoniae* (Dicty Stock Center DBS0349837) were grown in SM broth (55.51 mM glucose,1% proteose peptone, 0.1% yeast extract, 4.15 mM MgSO_4_, 13.96 mM KH_2_PO_4_, 3.44 mM K_2_HPO_4_, pH 6.2) supplemented with Ampicillin [100 µg/mL] on the benchtop (∼22°C). The next day, bacterial cultures were washed 3 times by repeated centrifugation at 3200g for 5 minutes and resuspension in fresh SM broth. Following washes, the culture volume was adjusted with SM broth to obtain an OD_600_ of 1. Amoeba cultures of the AX2 wild type or *tirBCD* single, double, and/or triple mutant amoebae were collected from confluent stationary cell culture flasks and counted on a hemocytometer. Amoeba density was adjusted to 1e3 cells/mL by serial dilutions in SM broth. 15 total amoeba cells were mixed with 600 µL of washed OD_600_=1 *K. pneumoniae* culture and spread onto dried SM/5 agar plates with sterile glass beads and allowed to dry fully. Plates were incubated for 3-5 days at 22°C and imaged with an Amersham Imager 600 on GFP settings. Plaque number and diameter were quantified by ImageJ.

### Amoeba fruiting body development

To test for any defects in the development of fruiting bodies, amoeba were grown up in t25 flasks at 22°C without shaking, and harvested at confluence. Amoeba were washed and concentrated in SorC before being plating on SM/5 agar plates with 0.5% charcoal at a density of 2.6e^6^ cells/cm^2^. We let them grow overnight and imaged them at 16hr, 20hr, and 24hrs.

### Infections with *Legionella pneumophila* and gentamicin protection assay

Amoeba mutant interactions with bacterial pathogens were tested by infecting cells with *Legionella pneumophila* and assessing bacterial intracellular replication and uptake. Amoeba were grown to confluence in t25 flasks at 22°C without shaking. Cells were collected from stationary flasks, washed 2 times with LoFlo (low fluorescence media, Formedium #L6001, pH 6.9) by centrifugation at 500g for 5 minutes and density was adjusted to 1.0×10^6^ cells/mL. 100 uL of amoeba were seeded into wells of a 96 well plate. Amoeba were incubated at 22°C while the bacteria were prepared (∼1 hour). A luminescent strain of KS79 containing the *Photorhabdus luminescens* luxCDABE operon was generated via triparental mating with KS79 *Legionella pneumophila, E. coli* strain TL154, and *E. coli* strain TL155 as detailed in Coers et. al. ^67^. A luminescent strain of KS79 *Legionella pneumophila* was grown overnight in AYE (54.88 mM ACES, 1% yeast extract, 2.28 mM L-cysteine hydrochloride, 611.48 mM FeSO_4_*7H_2_O, pH 6.9) at 37°C, with double orbital shaking in a Biotek Cytation 5 plate reader, measuring OD^600^ every 15 minutes. After ∼15 hours of growth, the bacteria were diluted in LoFlo media to a density of 2.5e7 CFU/mL. Four microliters of these cells were used to infect each amoeba strain. To synchronize the infections and maximize bacteria/amoeba cell contact, we centrifuged the samples at 220g for 2 minutes. The samples were then incubated at 25°C for 2 hours to allow time for the amoeba to phagocytose bacteria into the cells. After 2 hours, the adhered amoeba cells were washed 2x in LoFlo to remove extracellular bacteria. The 96-well plate was then placed in a SpectraMax iD3 luminometer and the luminescence of the bacteria was measured over the course of 72 hours with measurements every 30 minutes.

To quantify *L. pneumophila* uptake into amoeba cells, we performed a gentamicin protection assay. Both *Legionella pneumophila* and amoeba were grown and washed as for a regular *Legionella pneumophila* infection. After washing two hours post infection, cells were further washed in 100ug/mL Gentamicin in LoFlo media. After two washes, cells were incubated in 100 uL of 100ug/mL Gentamicin-Loflo for 45 minutes at 25°C. Next, cells were washed twice in 100 uL LoFlo. Amoeba cells were then lysed with 0.1% saponin in LoFlo at 37°C for 10 minutes. Cells were homogenized via vigorous pipetting. Lysates were then serial diluted in 1xPBS and plated on BCYE plates (54.88 mM ACES, 1% yeast extract, 166.53 mM activated charcoal, 1.5% agar, 2.28 mM L-cysteine hydrochloride, 618.81 mM Fe(NO_3_)_3_*9H_2_0, pH 6.9). To calculate the number of intracellular bacteria, we counted the number of viable CFU after 3 days of growth on BCYE plates at 37°C.

### TIR overexpression in *Dictyostelium*

To avoid potential toxicity due to TirC overexpression, we used inducible promoters to express full length TirC and TirC E162A proteins in *Dictyostelium* using GoldenBraid vectors ^68^. The Goldenbraid process involves first domesticating genes of interest, then assembling transcriptional units which are: regulatory elements (promoters and terminators), the gene of interest, flexible linkers, and protein tags into an Alpha-level vector. To express multiple transcriptional units from one vector, Alpha assemblies are then combined into Omega-level vectors. In place of traditional domestication, made-to-order plasmids containing full-length WT and E162A *tirC* sequences including appropriate 4bp grammar and overhangs were synthesized into a pUC57 backbone (GenScript). For the ΔNT TirC constructs, we used primers Supp. table 5) to amplify the *tirC* locus from AX4 gDNA to obtain the WT sequence. For the TirC E162A constructs, PCR used as a template a E162A made-to-order plasmid with appropriate 4bp grammar and overhangs for domestication. Then the ΔNT *tirC* amplicons were domesticated in the pUDP2 vector through Golden Gate assembly with BsmBI. Domestication assemblies were sequence verified before proceeding to the next cloning step. Next, we constructed WT and E162A TirC transcriptional units through Golden Gate assembly with BsaI (Alpha assembly). TirC transcriptional units included the following: a Tetracycline Response Element (TRE) as the promoter (amplified from pMB38 ^69^ [Dicty stock center plasmid ID: 45] and domesticated in pUDP2), the *tirC* sequences, flexible C terminal linkers, mNeon Green tags, and the *act8* terminator sequence in the a2 alpha level vector. For the GFP control vectors, we made transcriptional units consisting of the *act15* promoter sequence or the TRE, superfolder GFP ORF, and the act8 terminator in the a2 level vector. The Golden Gate product was transformed into NEB® Stable Competent E. coli (New England Biolabs,C3040H). Alpha assemblies were sequence verified before proceeding to the next cloning step. Extrachromosomal expression vectors were constructed through Golden Gate assembly with BsmBI (Omega assembly). We used the a1 transcriptional unit DdExChr, which contains several genes from the *D. discoideum* plasmid Ddp1 to allow for extrachromosomal plasmid maintenance and integrated this transcriptional unit with our TirC and GFP a2 transcriptional units onto the Omega-level vector O1H. The Golden Gate product was transformed into NEB® Stable Competent E. coli (New England Biolabs,C3040H).

Once the final vectors were sequence validated, we introduced 15 µg of each omega vector into 1.0×10^7^ MB35 (Dicty stock center DBS0236537, AX2 expressing Tet-OFF transactivator ^70^) cells via electroporation (as described in *Generating TIR deletion mutants in Dictyostelium amoebae* methods), and 24h later added 25 µg/mL Hygromycin B to select for transformed cells and 10 µg/mL doxycycline to prevent early *tirC*-mNeonGreen expression. We harvested polyclonal populations of transformants after ∼10 days post selection for functional assays and cell stocks. During functional assays, we observed protein expression ∼24 hours after Doxycycline removal.

### Fluorescence microscopy of amoeba cells and image analysis

To examine the localization and oligomerization of the TirC-mNeonGreen proteins, induced cells were washed in LoFlo supplemented with 25 µg/mL Hygromycin B, placed into Ibidi glass bottom cell chamber slides (Ibidi USA 80807), incubated for ∼1 hour at 22°C to allow cells to adhere, and observed with fluorescence microscopy. For Nicotinamide supplementation experiments, LoFlo medium was replaced from adhered cells with LoFlo supplemented with 20mM Nicotinamide and 25 µg/mL Hygromycin B. Epifluorescent images were taken on Nikon eclipse TE2000U and, respectively using 63x oil objectives. GFP and mNeonGreen fluorophores were excited with the 488 nm laser at 1% laser power and 50 millisecond exposure and emission was collected with a 500-530nm bandwidth emission filter. Brightfield exposure was set to 5 milliseconds. Images taken on Nikon eclipse TE2000U were processed in Fiji (ImageJ, v1.54p). For green fluorescent cell quantification, background fluorescence was normalized using Rolling Ball Background Subtraction and integrated density of cells was measured in the green channel. From this value, we calculated the corrected total cell fluorescence (CTCF) of cells and set an empirical threshold determined by cellular autofluorescence in non-GFP expressing cells. The percentage of green cells was calculated as the percentage of cells that had a CTCF greater than our threshold, divided by total cells.

## Acknowledgements

We thank Philip Kranzusch and members of the Levin lab for their feedback and Shriram Venkatesan and Jacob Jensen for assistance with data processing. Work in the Halfmann lab was funded by the Stowers Institute for Medical Research. Research conducted by Jocelyn Leon Padilla and Benjamin R. Morehouse was supported by the Institute of General Medical Sciences of the National Institutes of Health under Award Number R35GM157311. Research conducted by Edward Culbertson, Emily Cruz-Lorenzo, James Drurey, Neha Garlapti, Harshitha Gompa, and Tera Levin was supported by the Institute of General Medical Sciences of the National Institutes of Health under Award Number R35GM150681. This research was supported in part by resources from the University of Pittsburgh Center for Research Computing, RRID:SCR_022735. Specifically, this work used the HTC cluster, which is supported by NIH award number S10OD028483.

## Supplemental Figures, Tables, and Movies

*Legend:* Supp. Table. 1: **TIR domain containing proteins from phylogenetic analysis.**

A table displaying information on all proteins in the phylogenetic analysis (Fig. 1). Data includes the sequence name, full protein amino acid sequence, the domains present, the clade in the phylogenetic tree of Fig. 1, taxonomic information, and which domain of life the sequence is from. Domain information is displayed as <Name of the sequence>, <total length of protein>, <start of domain> | <end of domain> | <domain name> | <domain Pfam ID>. Multiple domains are separated by a comma.

*Legend:* Supp. Table. 2: Neighboring genes in the *tirB, tirC,* and *tirD* loci.

The neighboring genes (3 upstream, 3 downstream) around *tirB, tirC,* and *tirD* in the *Dictyostelium discoideum* AX2 genome are identified by the TIR locus, the name of the gene, the position of the gene relative to the nearby TIR gene, the gene annotation, and gene symbol. We searched both the genomic nucleotide sequences as well as the exonic sequences with nBLAST and display the top hit result in the table. Because we excluded all Dictyostelia, many of these genes yielded no BLAST hits (labeled as “No hit found”). We next translated all of these nucleotide sequences into amino acid sequences and used Hmmscan to determine what domains are present and used Jackhmmer to look for homology to other proteins. From Jackhmmer, we are reporting the taxonomic classification (at the domain level) of the significant hits. We then show the nucleotide sequences (both the genomic sequence and the exon-only sequence). Finally, we display the protein sequence.

*Legend:* Supp. Table. 3: **Strains used in Study**

*Legend:* Supp. Table. 4: **Plasmids used in Study**

*Legend:* Supp. Table. 5: **Primers used in Study**

*Legend:* **S6 File: TIR Atlas Tree, MUSCLE, 20%Trim**

Newick file of maximum likelihood phylogenetic tree of TIRs generated via MUSCLE alignment with TrimAL (-gt 0.2) with IQtree (v3.0.1, -m MFP, -bb 1000). Newick file is used in Figs 1 and Supp. Fig. 2. Node support calculated from ultrafast Bootstraps.

*Legend:* **S7 File: TIR Atlas Tree, MUSCLE, 10%Trim**

Newick file of maximum likelihood phylogenetic tree of TIRs generated via MUSCLE alignment with TrimAL (-gt 0.1) with IQtree (v3.0.1, -m MFP, -bb 1000). Newick file is used in Supp. Fig. 2. Node support calculated from ultrafast Bootstraps.

*Legend:* **S8 File: TIR Atlas Tree, MAFFT, 20%Trim**

Newick file of maximum likelihood phylogenetic tree of TIRs generated via MAFFT alignment with TrimAL (-gt 0.2) with IQtree (v3.0.1, -m MFP, -bb 1000). Newick file is used in Supp. Fig. 2. Node support calculated from ultrafast Bootstraps.

*Legend:* **S9 File: TIR Atlas Tree, MAFFT, 10%Trim**

Newick file of maximum likelihood phylogenetic tree of TIRs generated via MAFFT alignment with TrimAL (-gt 0.1) with IQtree (v3.0.1, -m MFP, -bb 1000). Newick file is used in Supp. Fig. 2. Node support calculated from ultrafast Bootstraps.

*Legend:* **S10 File: Clade 1 Tree**

Newick file of maximum likelihood phylogenetic tree of Clade 1 TIRs generated via MUSCLE alignment with TrimAL (-gt 0.05) with IQtree (v3.0.1, -m MFP, -bb 1000). Newick file is used in Supp. Fig. 3. Node support calculated from ultrafast Bootstraps.

*Legend:* **S11 File: Clade 4 Tree**

Newick file of maximum likelihood phylogenetic tree of Clade 4 TIRs generated via MUSCLE alignment with TrimAL (-gt 0.05) with IQtree (v3.0.1, -m MFP, -bb 1000). Newick file is used in Supp. Fig. 3. Node support calculated from ultrafast Bootstraps.

*Legend:* **S12 File: Clade 6 Tree**

Newick file of maximum likelihood phylogenetic tree of Clade 6 TIRs generated via MUSCLE alignment with TrimAL (-gt 0.05) with IQtree (v3.0.1, -m MFP, -bb 1000). Newick file is used in Figs 1. Node support calculated from ultrafast Bootstraps.

*Legend:* **S13 File: Clade 8 Tree**

Newick file of maximum likelihood phylogenetic tree of Clade 8 TIRs generated via MUSCLE alignment with TrimAL (-gt 0.05) with IQtree (v3.0.1, -m MFP, -bb 1000). Newick file is used in Supp. Fig. 3. Node support calculated from ultrafast Bootstraps.

*Legend*: Supp. Mov. 1: ***D. discoideum* expressing full length WT tirC**

Fluorescent microscopy time lapse of *D. discoideum* expressing full length WT tirC-mNeonGreen. The time lapse covers 16 hours with a 10 minute frame interval. The time stamp in the top right corner displays times in an hour:minute format.

*Legend*: Supp. Mov. 2: ***D. discoideum* expressing tirC ΔNT**

Fluorescent microscopy time lapse of D. discoideum expressing tirC ΔNT-mNeonGreen. The time lapse covers 18 hours with a 10 minute frame interval. The time stamp in the top right corner displays times in an hour:minute format. Boxes highlight mNeonGreen-positive cells undergoing cell lysis.

**Figure S1.**
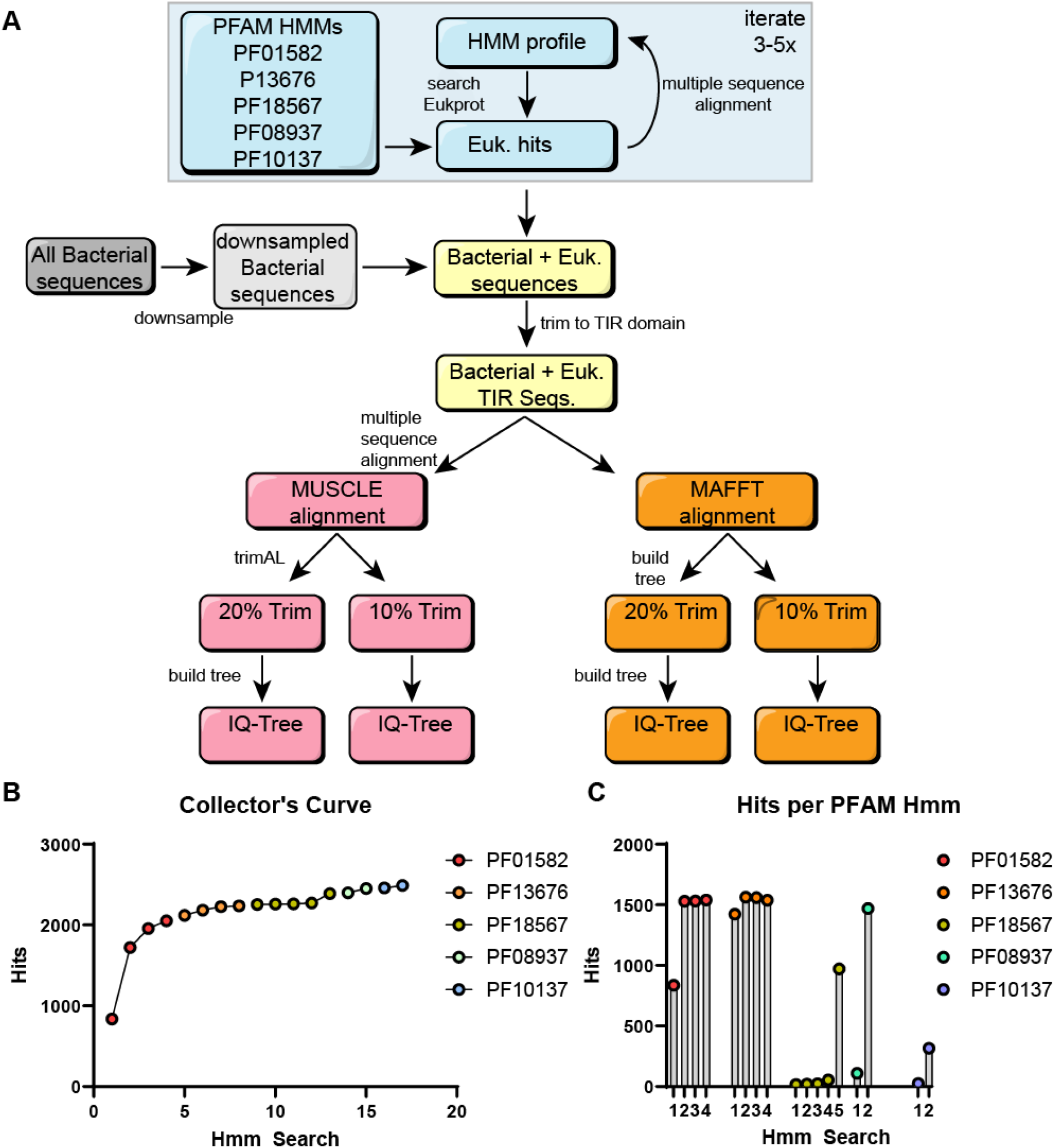
TIR domain search across multiple PFAM HMMs. **A.** Schematic of search strategy employed to find diverse TIR domains across the eukaryotic tree of life. **B.** Collector’s curve of searches (left) and **C.** the total number of hits per Pfam Hmm search.

**Figure S2.**
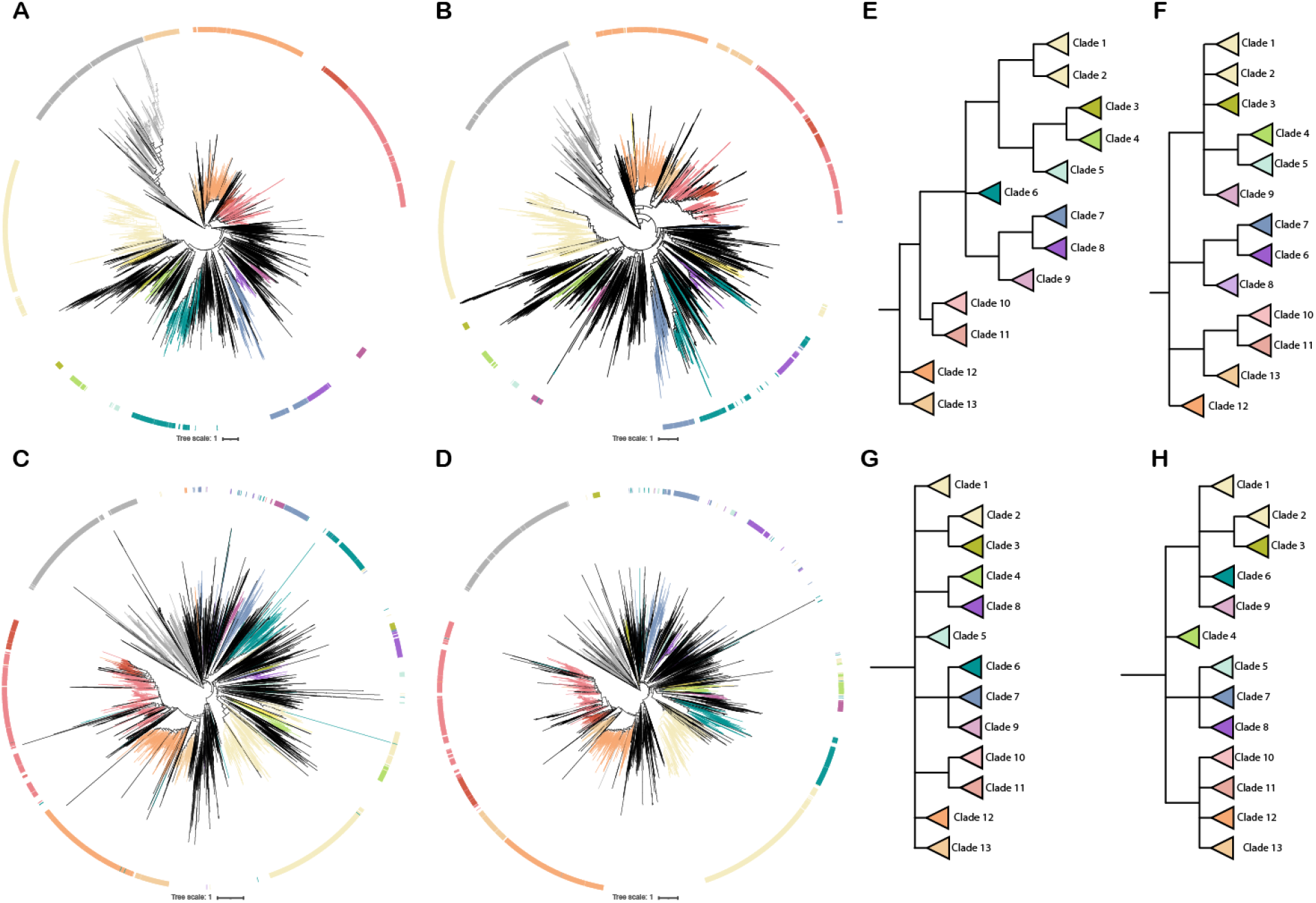
TIR domain clades are robust across alignment parameters. Four phylogenetic trees generated by IQtree from varied alignments, showing clades supported across all trees and a schematic of the clade topologies. **A.** Alignment generated via MUSCLE with a 20% trim. **B.** Alignment generated via MUSCLE with a 10% trim. **C.** Alignment generated via MAFFT with a 20% trim. **D.** Alignment generated via Mafft with a 10% trim. **E-H,** Collapsed clades from the four IQtrees (**E**-Muscle 20% trim, **F**-Muscle 10% trim, **G**-MAFFT 20% trim, **H**-MAFFT 10% trim) with clades colored as in Fig. 1. Only nodes with 50 or greater bootstrap values are shown, unsupported nodes are collapsed to polytomies.

**Figure S3.**
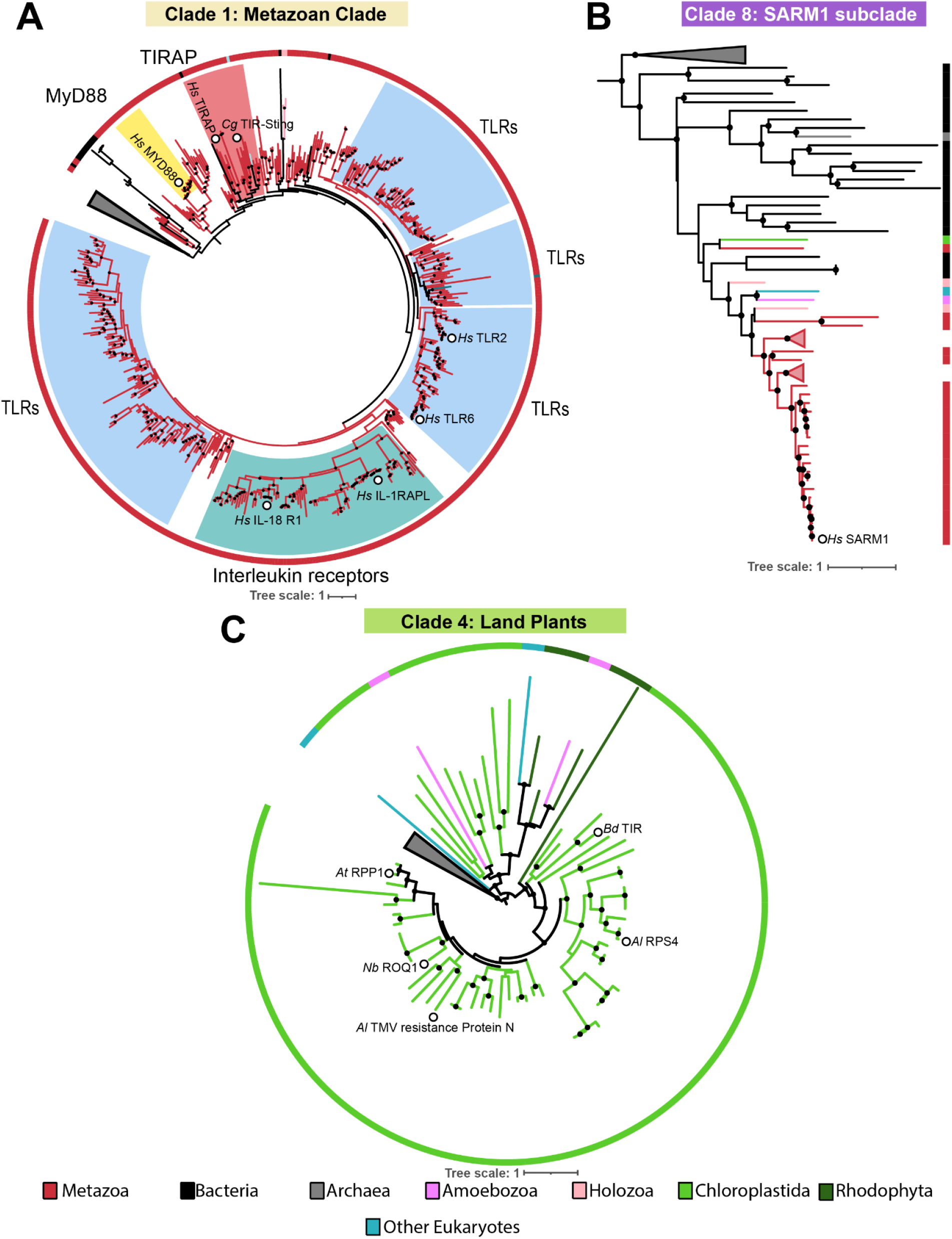
TIRs family members in clades 1 & 4. **A.** A circular maximum likelihood phylogenetic tree generated by IQtree of TIR domains from Clade 1. The majority of sequences in this clade were Metazoan TIRs. Colored wedges designate protein clades: MyD88 (yellow), TIRAP (red), TLRs (blue), Interleukin receptors (dark teal). **B.** Maximum likelihood phylogenetic tree generated by IQtree of TIR domains from Clade 8. Clade 8 was a mainly bacterial clade into which a subset of Metazoan TIRs, notably SARM1, clustered. **C.** A circular maximum likelihood phylogenetic tree generated by IQtree of TIR domains from Clade 4. This clade contained all of the TIR domains from land plants (Embryophyta). White circles indicate notable TIR domain-containing proteins: (**AB**) *Hs* = *Homo sapiens,* (**C**) *At* = *Arabidopsis thaliana*, *Al* = *Arabidopsis lyrata, Bd* =*Brachypodium distachyon, Nb = Nicotiana bethamiana.* Both leaf color and strip color indicate the eukaryotic supergroup. Wedges represent collapsed parts of the phylogenetic trees. Trees are rooted on outgroup sequences (gray wedges). Ultrafast bootstrap values calculated by IQtree at all nodes with support >70 are shown as black dots.

**Figure S4.**
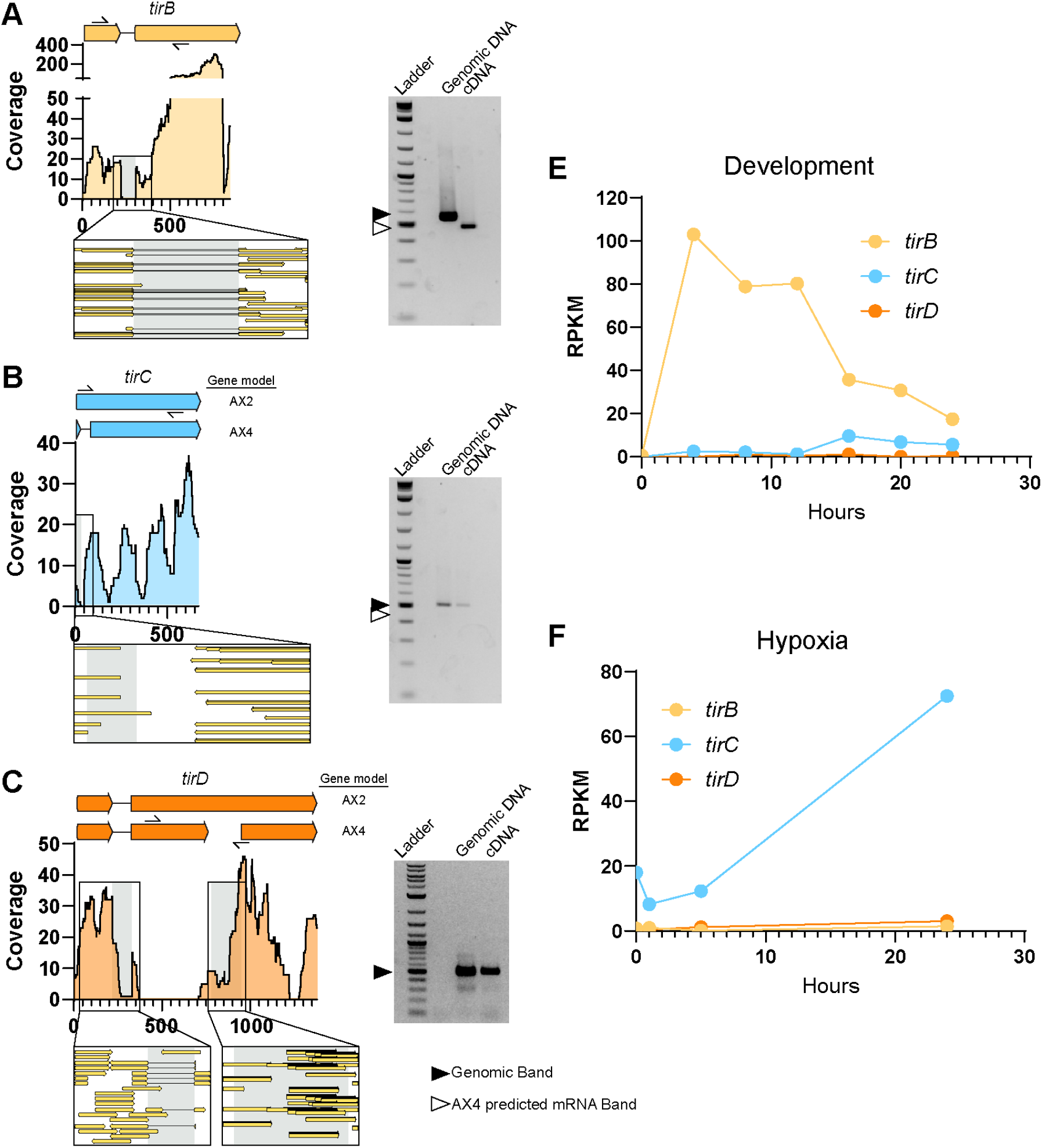
*tirBCD* gene models. **A-C.** Diagrams of predicted gene models for *tirB* (**A**), *tirC* (**B**), and *tirD* (**C**). For each gene, the AX2 and AX4 annotations are shown to scale. For *TirB* the two annotations agree and so a single schematic of the coding sequence is shown. For *TirD*, the AX4 annotation predicted that the gene was two genes (*DDB_G028737* and *DDB_G0287321*). Below each schematic is a plot showing the coverage via RNAseq at each position on the genomic locus. The focus box zooms in on the indicated area to show the alignment of individual reads to the genomic locus. The gray boxes indicate the predicted introns by both AX2 and AX4 and are displayed on both the coverage plots and the focus boxes. To the right of the RNAseq cover plots is an Agarose gel picture where the PCR products of the primers indicated on the gene locus schematics were used to amplify gDNA or cDNA. The black arrow indicates the expected product size when amplifying genomic DNA while the empty arrow shows the product size of amplified cDNA based on the AX4 genome annotation. For *TirD*, the reverse primer spans the AX4-predicted intron-exon boundary and so no product is predicted. **E.** Reads per Kilobase of transcript per Million reads mapped (RPKM) of *TirB, TirC,* and *TirD* (*DDB_G0287321*) from Parikh et. al. ^46^ over 24 hours of development on a filter. **F.** Reads per Kilobase of transcript per Million reads mapped (RPKM) of *TirB, TirC,* and *TirD* (*DDB_G0287321*) from Hesnard et. al. ^45^ over 24 hours of hypoxia exposure.

**Figure S5.**
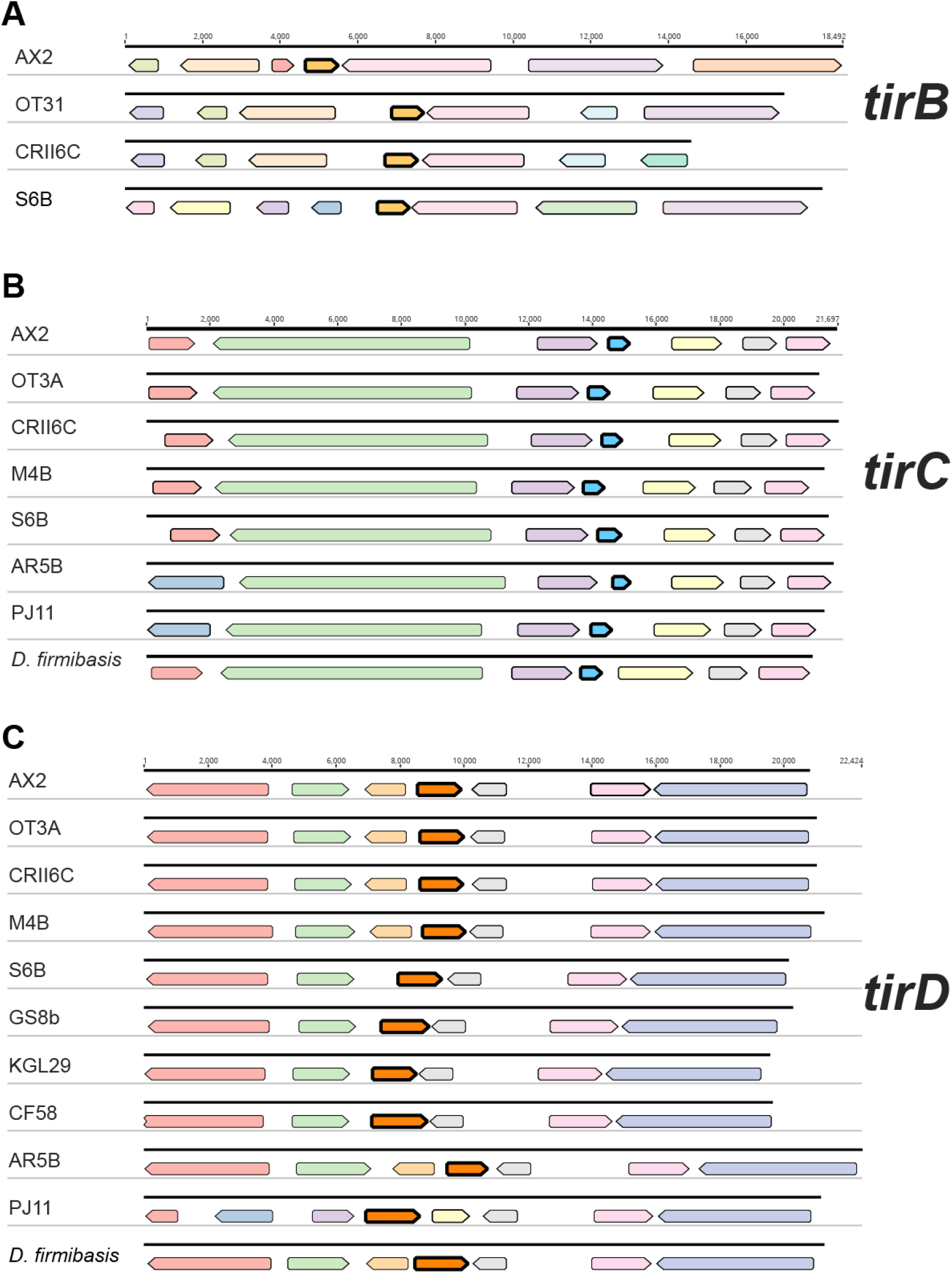
Syntenic distribution of genes at TirB, TirC, TirD genomic loci. Genetic synteny across Dicyostelium species at the (**A**) TirB, (**B**) TirC, and (**C**) TirD loci. Colored arrows represent homologous ORFs across the genomes, with strain identifiers listed on the left side. Genes of the same color within a given locus are homologous. Black outlined arrow represents the TIR-domain containing gene. Numbers above the loci show the nucleotide scale of the locus and genes.

**Figure S6.**
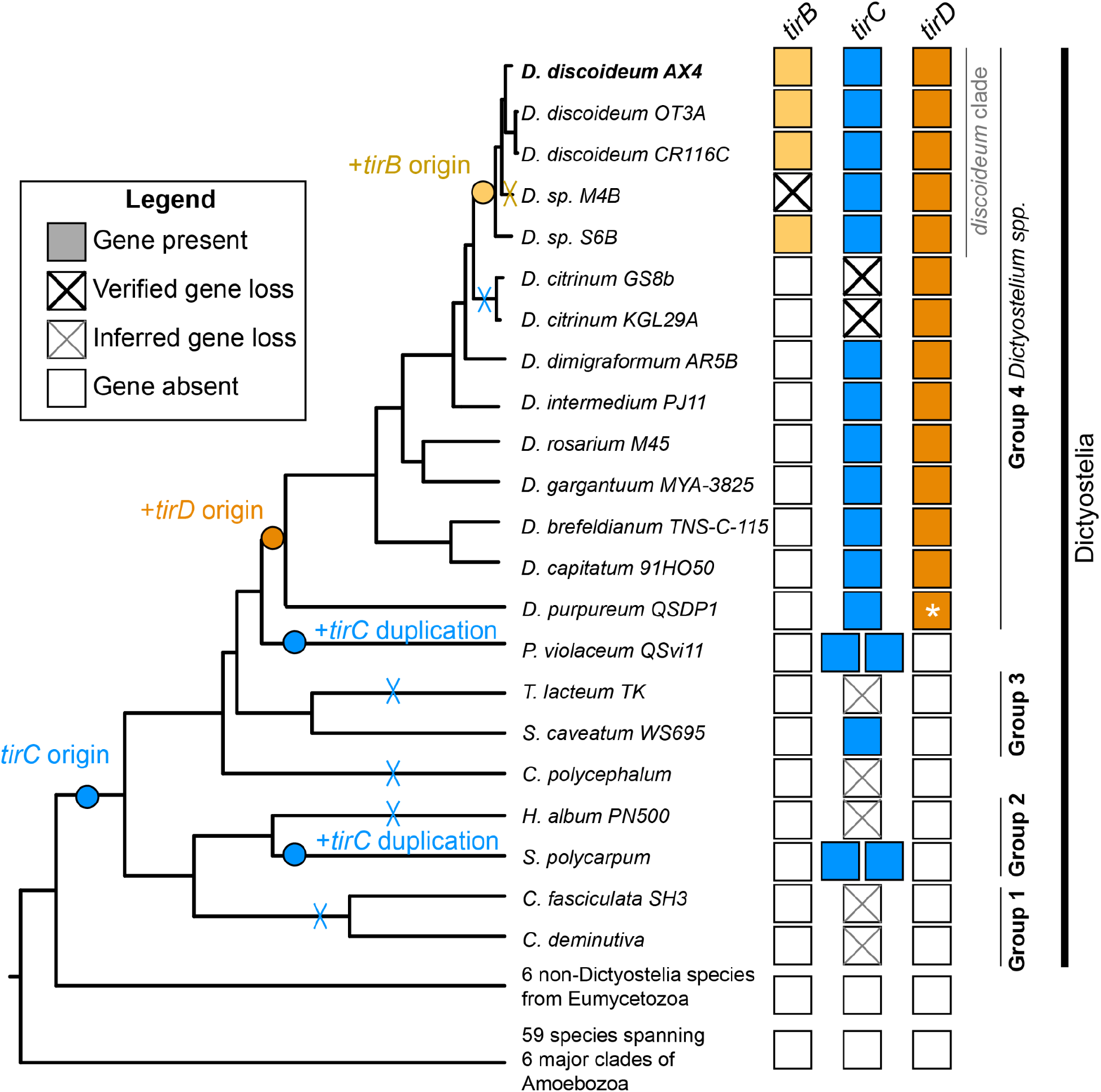
Origin and duplications of *tirBCD* genes within Dictyostelia. Colored boxes indicate species of cellular slime mold amoebae where *tirB*, *tirC*, and *tirD* genes were present, based on tBLASTn and TIR HMM searches of the amoeba genomes. All colored boxes were discovered by tBLASTn or both methods, except for *D. purpureum* TirD (*), which was identified only by a TIR HMM search. Black Xs mark instances of gene loss where we identified either pseudogenes or deletions in the syntenic locus. Gray Xs mark inferred losses based on the phylogeny, although these apparent absences could result from incomplete assemblies. Outgroup Amoebozoa and Eumycetozoa datasets searched included a combination of genomes and transcriptomes within Eukprot v3. Species tree is based on Schilde et. al. ^47^, branch lengths not to scale.

**Figure S7.**
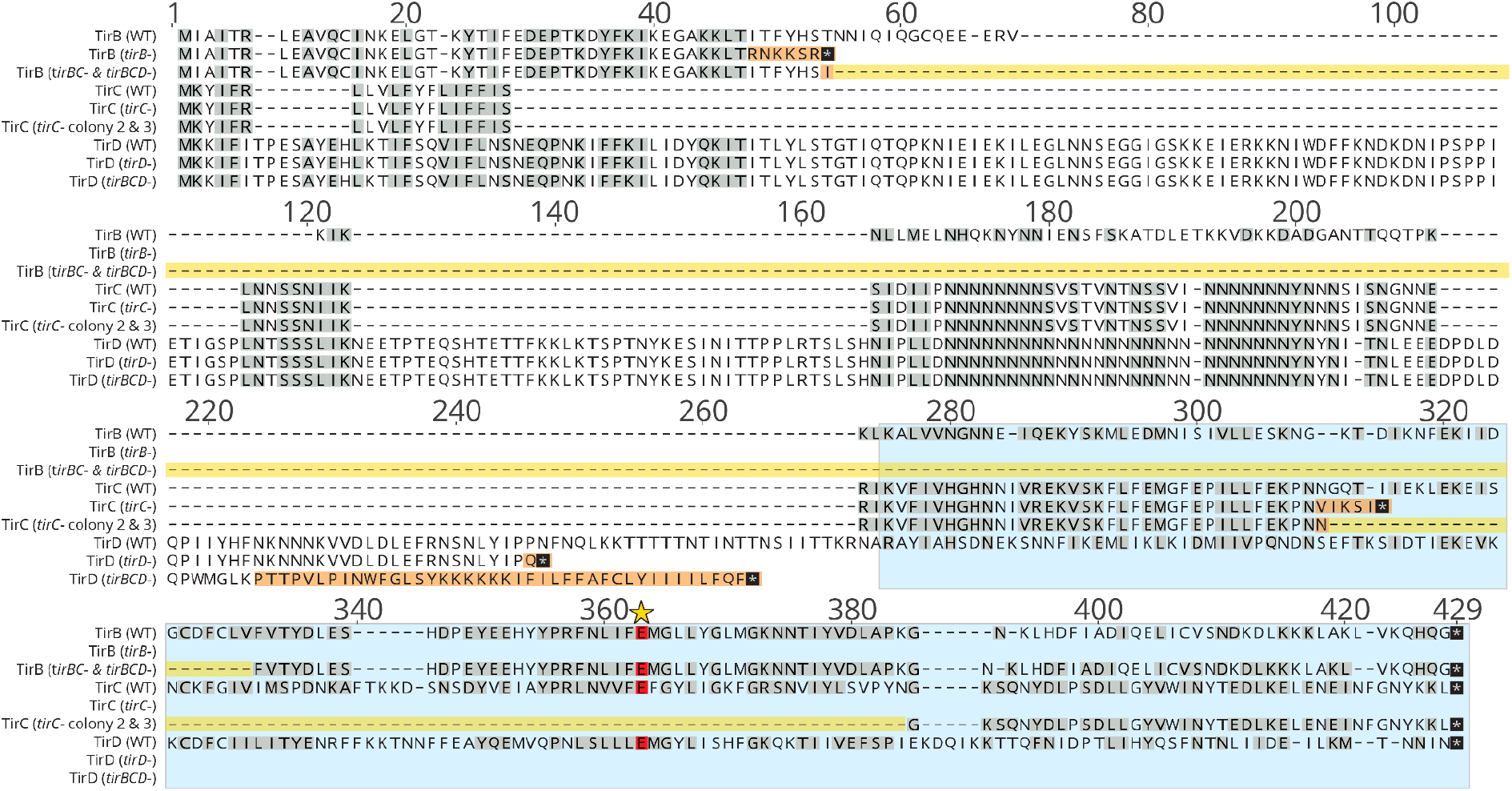
Amino acid alignment of TirB, TirC, and TirD and their mutants. Amino acid alignment of wild type TirB, TirC, and TirD and the predicted protein products made by genetically modified strains. For each protein, the wild type sequence is listed first, followed by predicted protein products of the mutant clones. The blue box indicates the TIR domain of the proteins. The orange boxes indicate frameshift induced mutations in the protein. The yellow box highlights the amino acids of the protein that have been deleted. Stop codons are highlighted with a black box. The catalytic glutamate is designated with a red box and the locus is marked with a yellow star.

**Figure S8.**
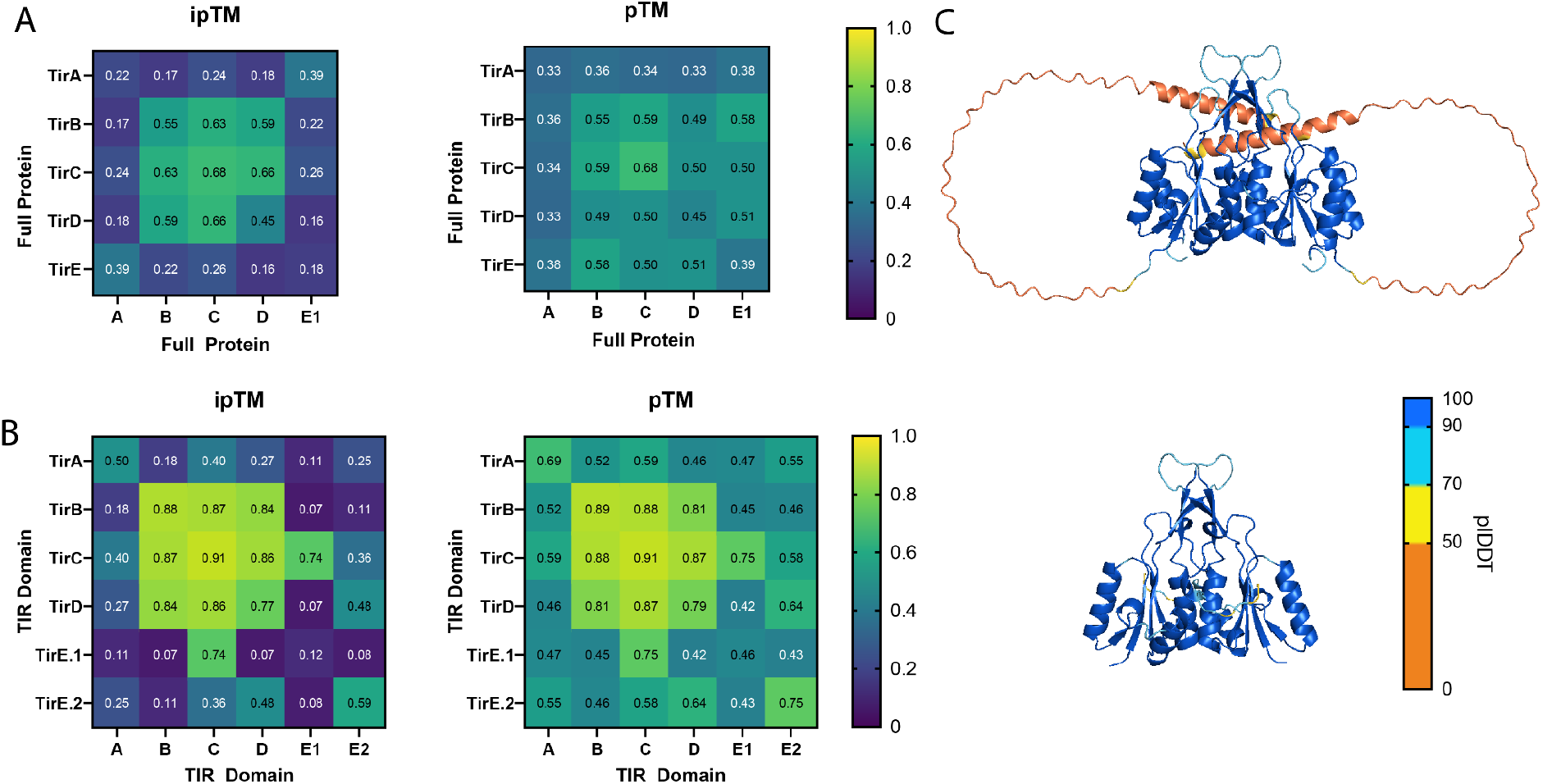
TirC is predicted to homodimerize. **A.** ipTM and pTM scores of AlphaFold Multimer predictions of *D. discoideum* TIR full protein dimers. **B.** ipTM and pTM scores of AlphaFold Multimer predictions of *D. discoideum* TIR domain dimers.**C.** AlphaFold models of full length (top) and TIR domain (bottom) TirC homodimers. Residues are colored by pIDDT score.

**Figure S9.**
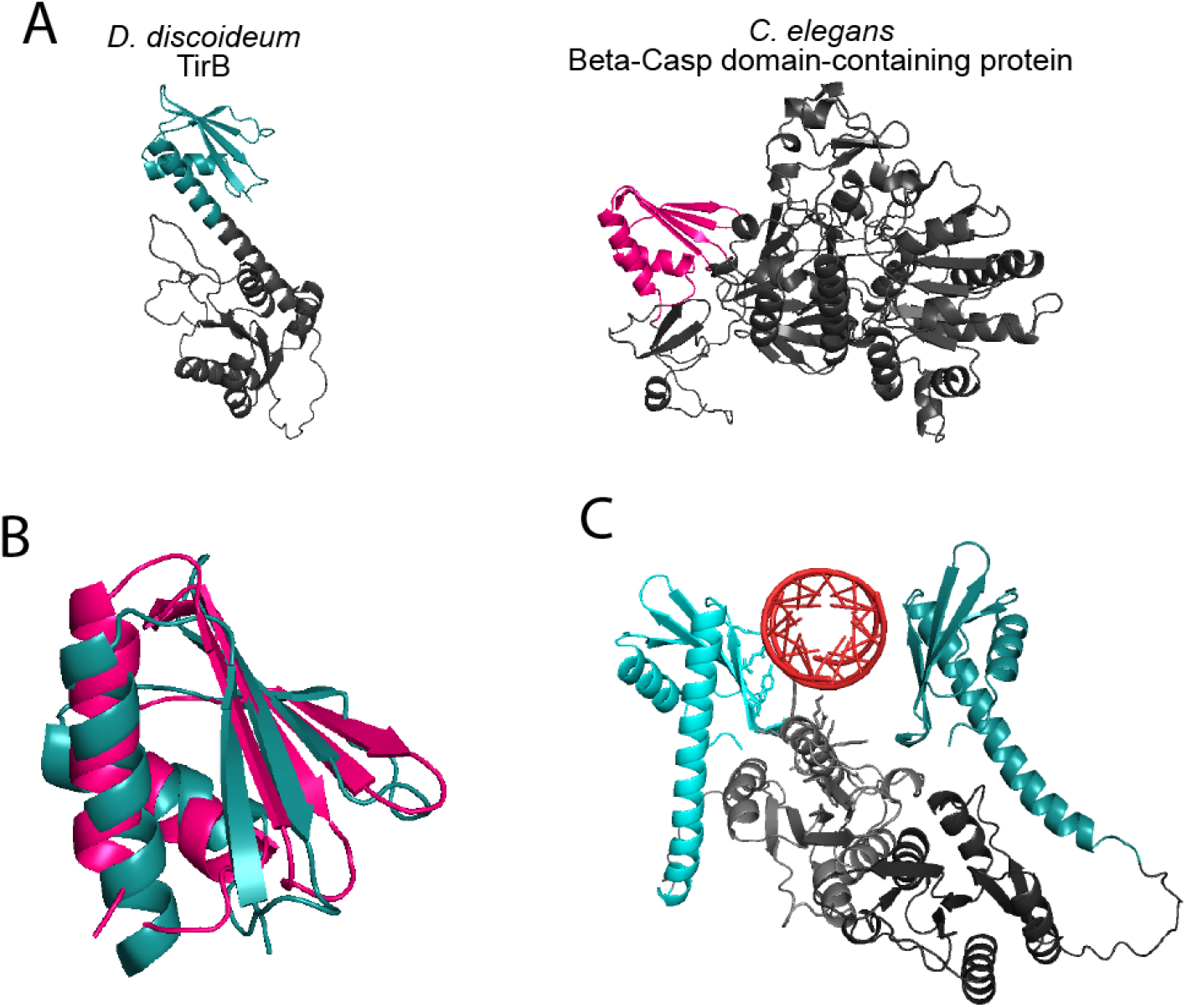
Predicted RNA binding in TirB’s N-terminus. **A.** AlphaFold predictions of *D. discoideum* TirB and *C. elegans* Beta-Casp domain-containing protein, the highest scoring FoldSeek hit for TirB’s N-terminus. The similar domains are colored, teal in TirB, and pink partial KH domain. **B.** Structural alignment of the KH domain of *C. elegans* Beta-Casp domain-containing protein and TirB’s N-terminus. The two structures aligned with an RMSD of 3.8 over 45 atoms. **C.** AlphaFold models of full length TirB-TirB homodimers binding a dsRNA molecule. dsRNA is shown in red, while the N-Termini of TirB are shown in shades of teal, and the TIR domain in gray. TirBs are differentiated by shading (protomer 1 is light, protomer 2 is dark).

**Figure S10.**
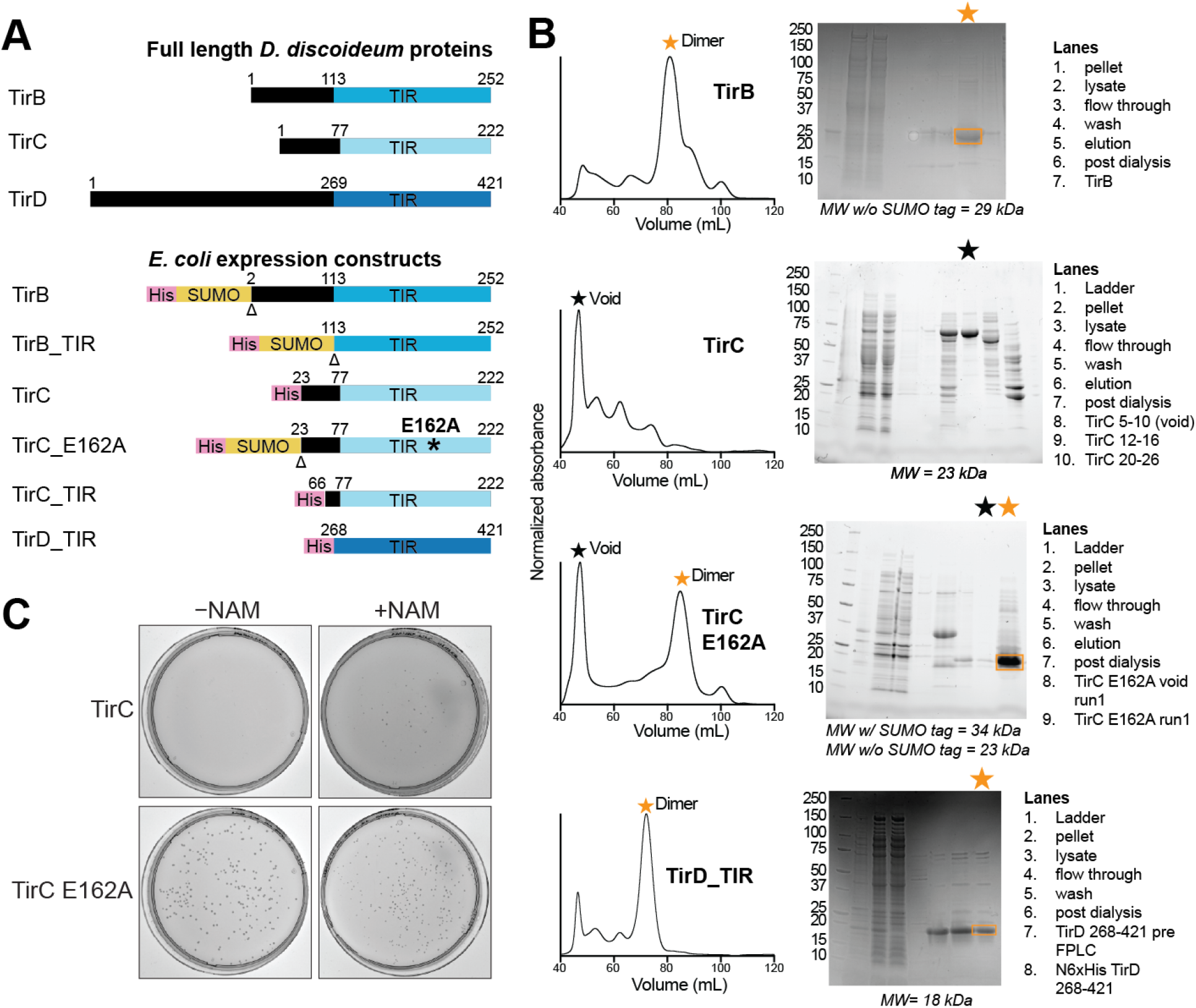
Purification of TirB, TirC, and TirD. **A.** Schematic of full length proteins and constructs. The SUMO2 tag was proteolytically removed before experimentation. His-tagged proteins had GS linkers between the tag and the protein. Numbers indicate the amino acid coordinates of TirB, C, and D proteins that were included. The TirC construct excluded the N-terminal hydrophobic alpha helix to facilitate protein expression and purification. * shows the location of the E162A point mutation. Δ indicates the SUMO2 cleavage site. **B.** Recombinant protein was purified via size exclusion chromatography (traces on left) and purity of fractions were determined by SDS-PAGE (gel images on right). Lanes are numbered from left to right. Stars indicate primary fractions considered for activity assays. Note that for WT TirC there was no obvious band on the gel and yet the void fraction contains measurable NADase activity (as shown in other figures). Contaminants near 20 kDa and 75 kDa are common E. coli proteins which bind nickel-resin with appreciable affinity under the conditions tested. **C.** TirC is toxic to E. coli, but partially rescued by extracellular nicotinamide. Images show colonies of bacteria on MDG agar plates in the absence of inducer, with toxicity due to leaky TirC expression from the T7 promoter. The E162A TirC mutant construct was not toxic.

**Figure S11.**
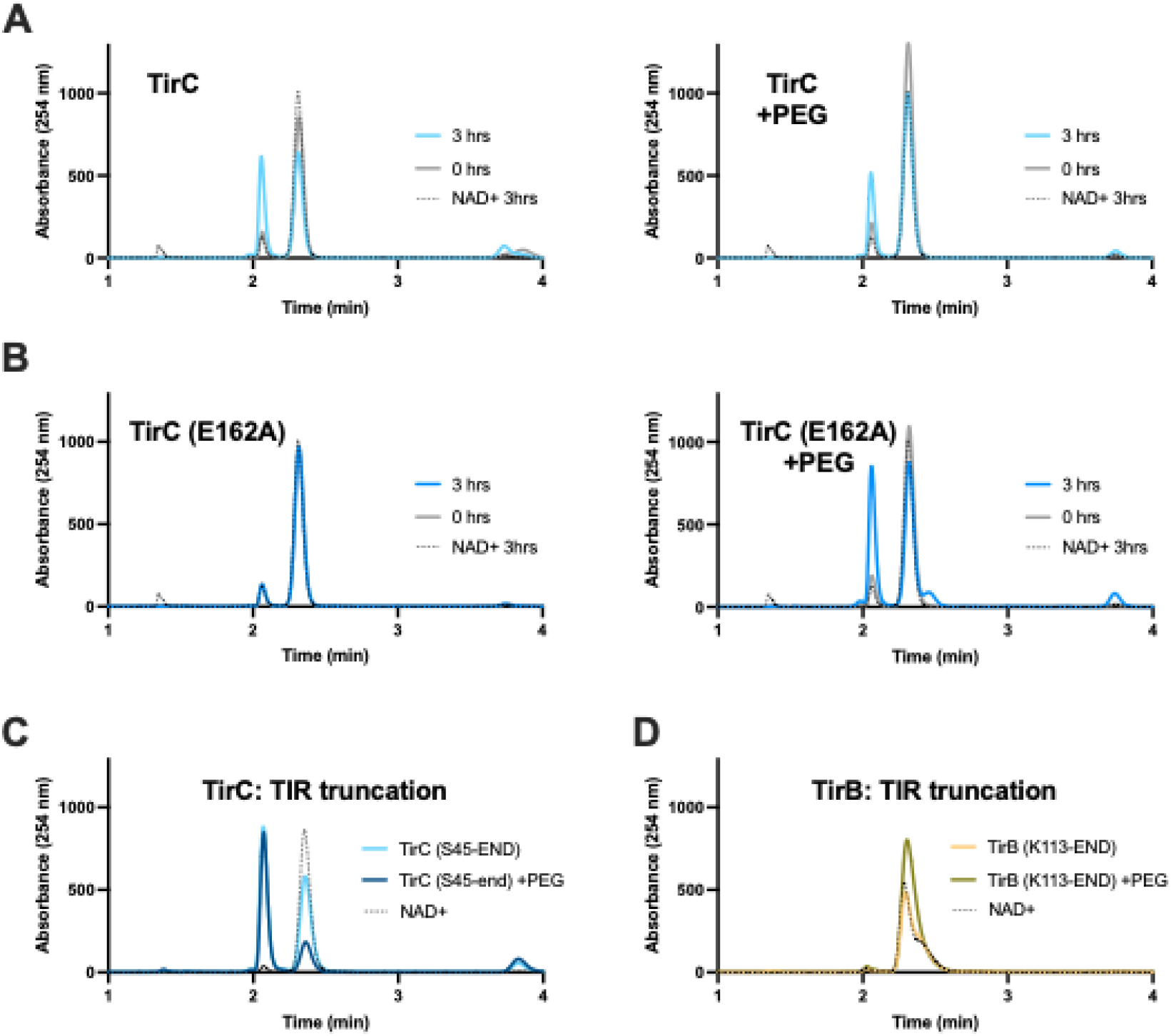
NAD^+^ processing by TirC and TirB mutants in the presence of PEG 8000. **A.** Time course of *in vitro* co-incubations of NAD^+^ and WT TirC in the presence of PEG 8000 as a molecular crowding agent. In both cases, WT TirC cleaves NAD^+^ to produce ADPR and NAM. Black dotted lines show the absorbance of NAD^+^ alone, following 3h incubation in the study conditions. Colored lines show traces after 0-3h co-incubation with protein. **B.** Similar time course using the TirC E162A mutant, which cleaves NAD^+^ only in the presence of PEG 8000. **C.** TirC truncation mutant (S45-END) that includes only the TIR domain processes NAD^+^ similar to TirC WT, with only a slight change in processing rate in the presence of PEG. **D.** Truncation mutant of TirB (K113-END) to the TIR domain does not show any NAD^+^ cleavage activity, even in the presence of PEG. Similar results were obtained for the TIR domain of TirD (shown in Fig 3).

**Figure S12.**
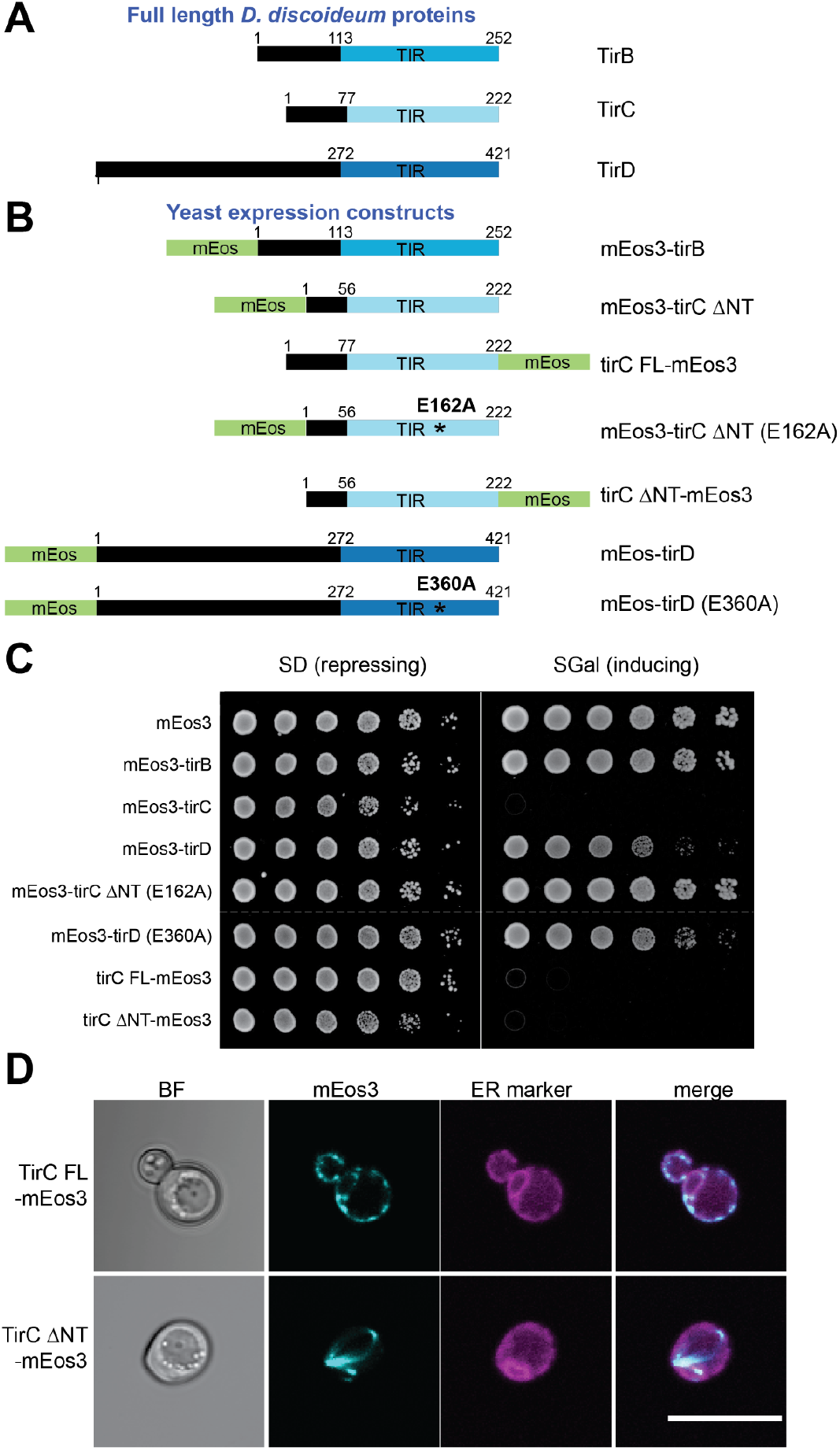
Expression of TirBCD in yeast. **A & B**. Schematic of *D. discoideum* TIR proteins expressed in *S. cerevisiae*. Numbers indicate the coordinates of TirB, C, and D proteins that were included. FL = full-length, ΔNT = N-terminal helix is missing, * shows the location of the putative catalytic glutamate. To facilitate DAmFRET and protein localization studies, the amoeba proteins were tagged with mEos fluorophores. **C**. Viability assay of yeast carrying the TirBCD expression constructs were serially diluted (5-fold) and spotted onto selective media either to repress (SD) or induce (Sgal) protein expression. **D.** Localization of TirC FL or ΔNT proteins in yeast expressing the mTagBFP2 ER marker. Cells were grown in excess nicotinamide to enhance TirC expression. TirC FL strongly co-localizes with the ER but TirC ΔNT does not, suggesting that the N-terminal helix facilitates membrane localization.

**Figure S13.**
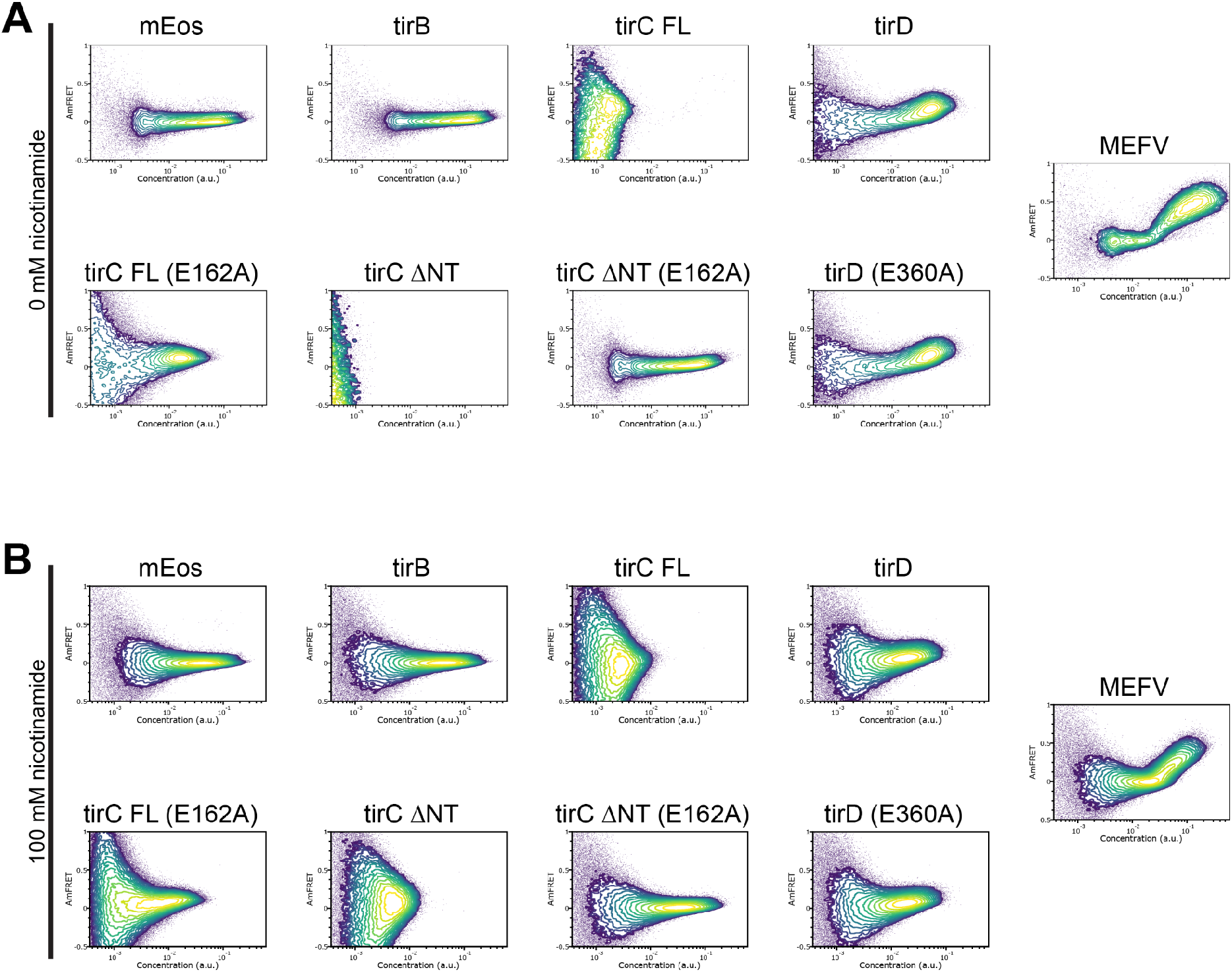
Self oligomerization of TIR proteins via DAmFRET. Representative DAmFRET data showing the propensity of various *D. discoideum* TIR constructs to oligomerize in yeast cells. mEos serves as a negative control, which fails to aggregate even at high concentrations, while MEFV is a positive control for aggregation and filamentation. Cells were grown without supplemental nicotinamide (**A**) and with 100 mM nicotinamide (**B**).

**Figure S14.**
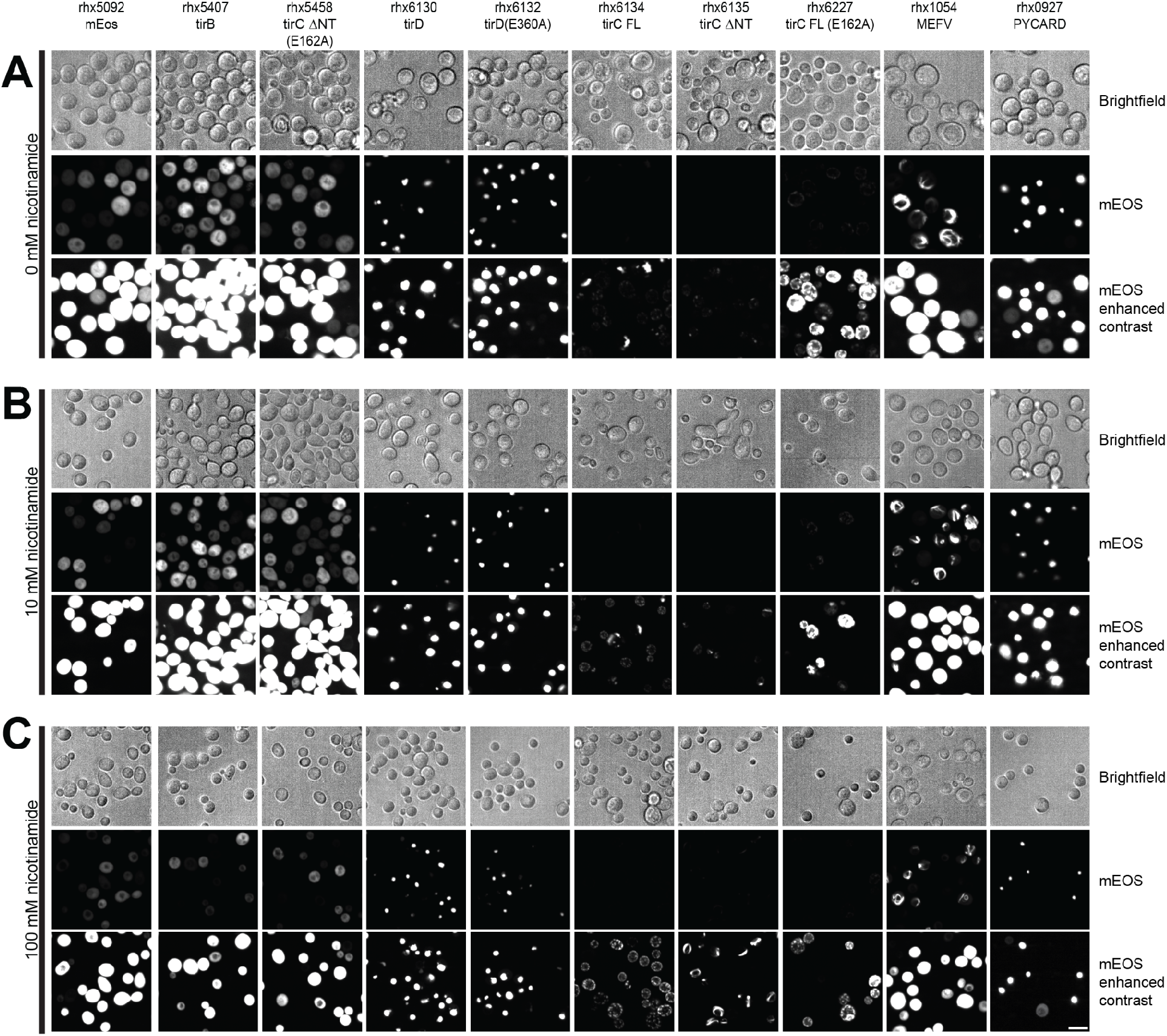
TIR construct localization in yeast. Microcopy images of yeast cells expressing various mEos3-tagged constructs. Images are from brightfield and the green (mEos3) channel. The mEOS channel is displayed twice with the contrast on the second image being enhanced to allow visualization of low mEos3 expression. Yeast cells were grown without supplemental nicotinamide (**A**), with 10 mM nicotinamide (**B**), and with 100 mM nicotinamide (**C**). Scale bar is 10 µm.

**Figure S15.**
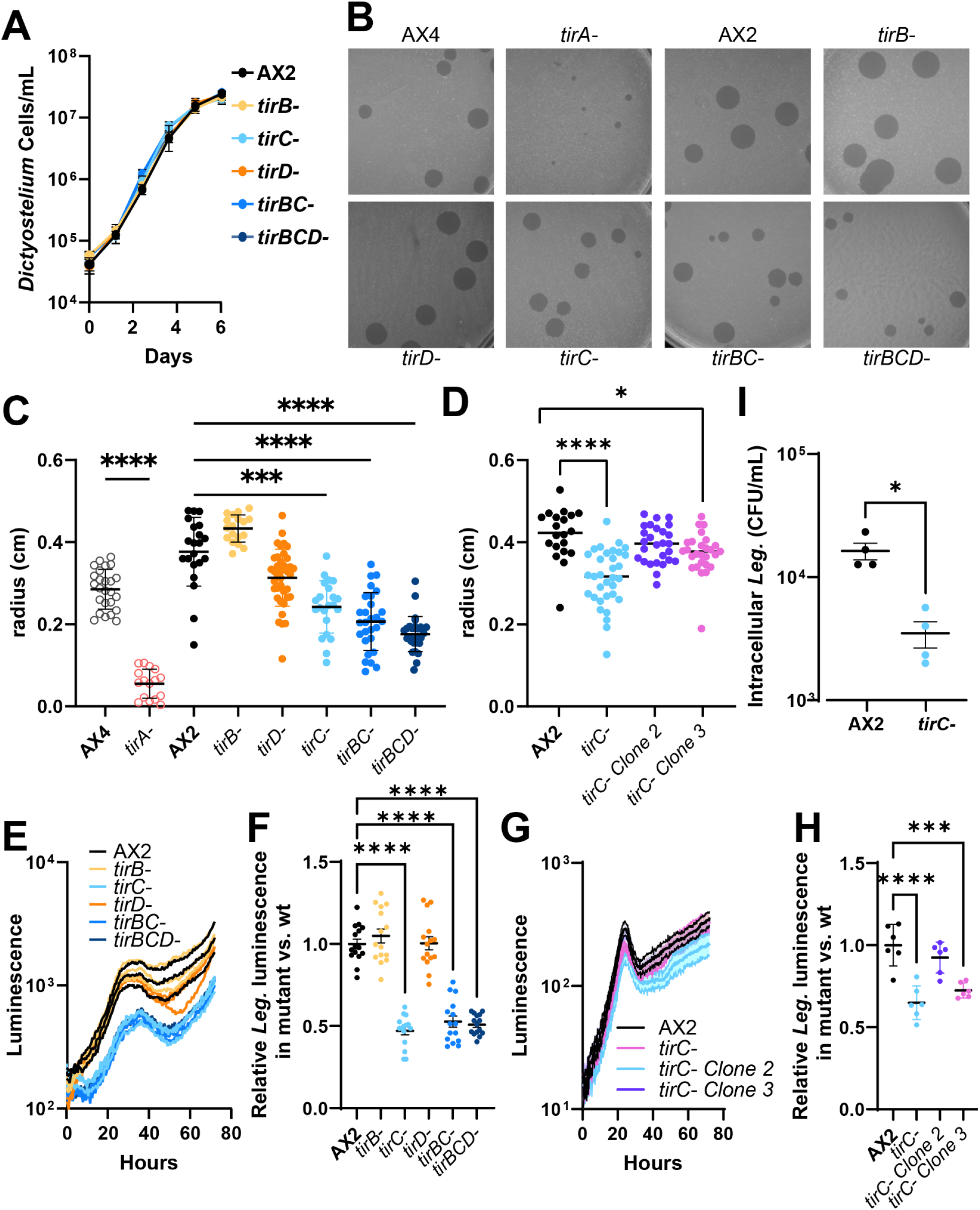
*tirC-* mutants are defective in bacterial uptake. **A.** Growth curve of wild type (AX2) and TIR mutant *D. discoideum* cultures in HL5 rich media. The TirBCD proteins are not essential for cell viability or growth. **B.** Representative photos of *D. discoideum* plaque sizes on a lawn of *K. pneumoniae* measured after four days of growth for all *D. discoideum* strains. **C-D.** Quantification of plaques across three replicate plates per genotype, measured after four days. Panel **C** shows plaque size across multiple genotypes while panel **D** shows plaque size variation amongst different *tirC-* clones. **E,G.** Curves of luminescent *Legionella pneumophila* (Leg) during infection of *D. discoideum* TIR mutants over 72 hours. *D. discoideum* cells were infected with Leg at an MOI of 10 and incubated at 22°C for three days. Experiment was carried out in triplicate wells with the luminescence curve of each well plotted. **F,H.** Comparison of the peak luminescence relative to wild-type reached by each strain between the 20-40 hours post infection window. Panel **E-F** shows Leg luminescence across multiple genotypes while panel **G-H** shows variation amongst different *tirC-* clones. **I.** Gentamicin protection assay during Leg infection of AX2 and *tirC-*. Error bars show standard deviations. Statistical significance for C,D, was determined by one-way ANOVA with Kruskal-Wallis posttest and statistical support for F,H was determined by one-way ANOVA with Kruskal-Wallis posttest; *p ≤ .05,***p ≤ .001, ****p ≤ .0001. Statistical significance for I was determined by two tailed student’s t-test; *p ≤ .05.

**Figure S16.**
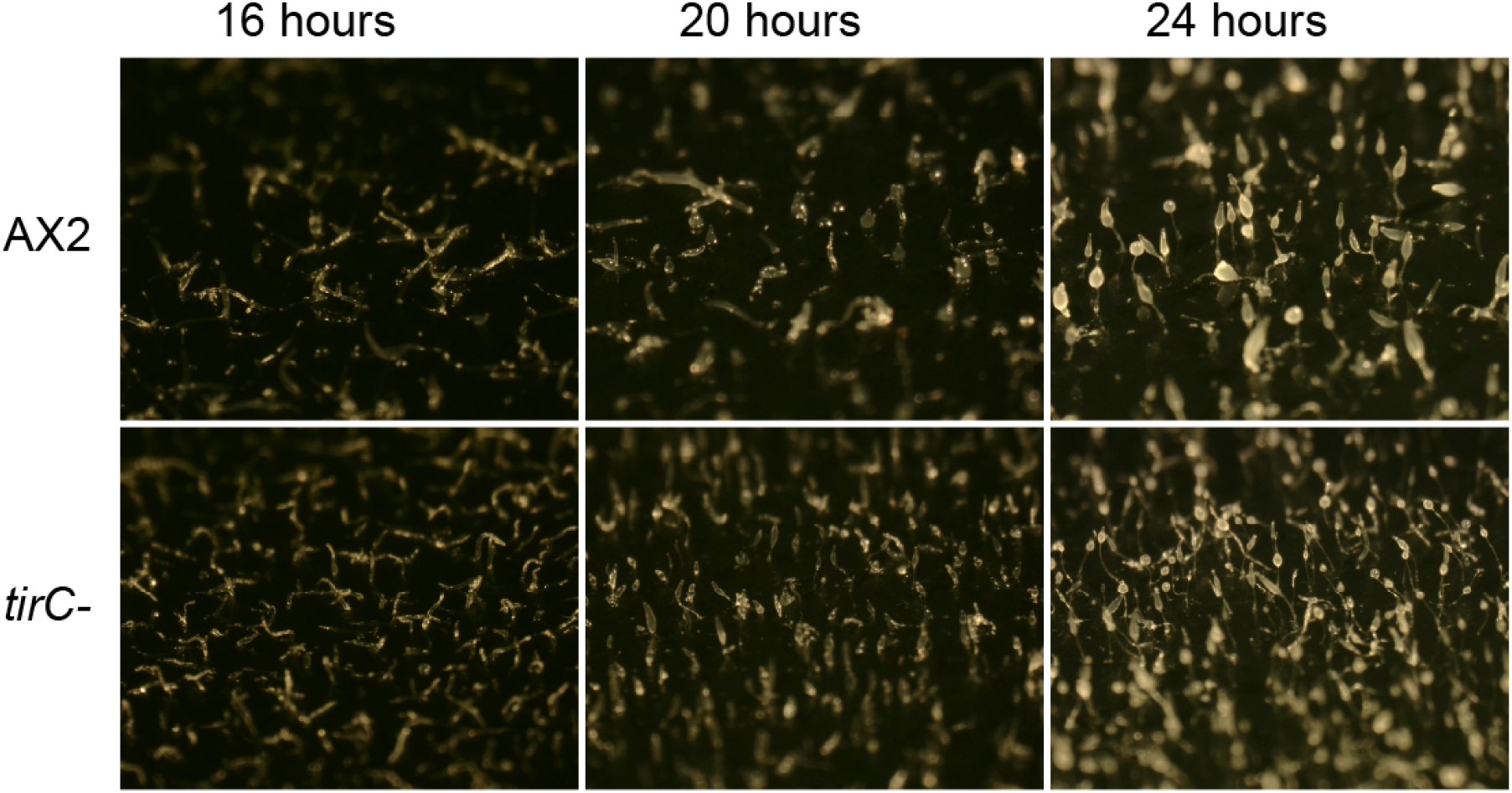
Development of fruiting bodies. Representative photographs of the development of fruiting bodies in D. discoideum wild type (AX2) and tirC- over 24 hours on SM/5 agar plates with 0.5% charcoal. D. discoideum cells aggregate into distinct morphological forms during the transition from unicellular to multicellular states. At 16 hours for both AX2 and tirC-, a mixture of slugs (horizontal rods) and fingers (vertical rods) are observed. At 20 hours, early culminants are visible, which are characterized by an asymmetrical main body that is smaller at the apical tip, held aloft by a thin stalk. At 24 hours, a mixture of mature fruiting bodies, with spherical sori atop thin stalks, and early culminants are visible for both strains.

**Figure S17.**
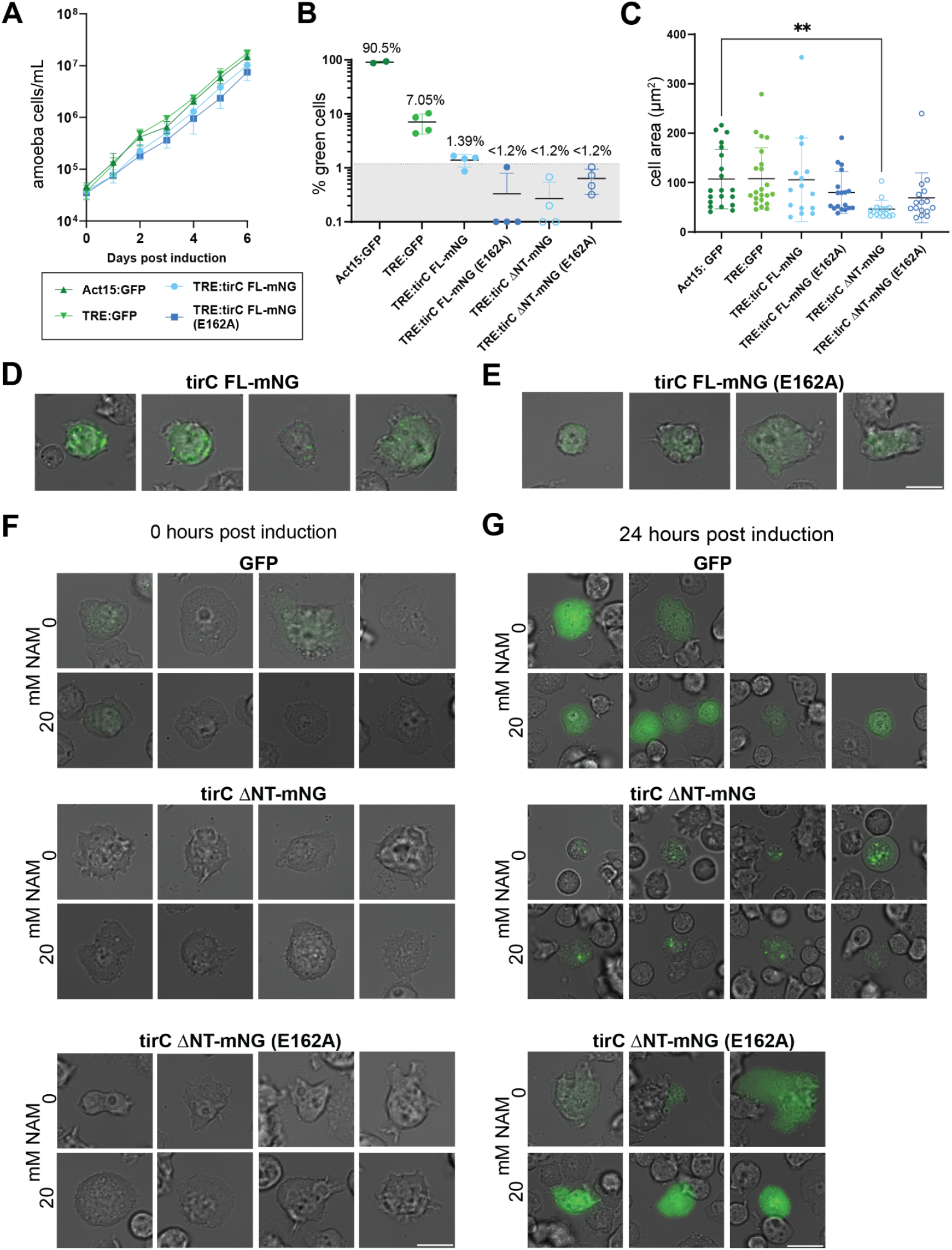
TirC overexpression in *D. discoideum*. **A.** Cell growth assay for amoebae expressing proteins under constitutive (Act15) or inducible (TRE) promoters. Doxycycline was removed from the cells at Day 0 to induce overexpression of polyclonal transformants expressing mNeonGreen-tagged (mNG) TirC or GFP controls. All overexpression cell lines grew at a similar rate. **B.** Quantification of green fluorescent cells in the polyclonal populations. Each dot represents at least 80 counted cells. Shaded area shows the approximate limit of detection, with symbols on the x-axis indicating fields of view that had no detectable green cells. **C.** Quantification of cross-sectional cell area for overexpression cell lines. Statistical significance was tested by a one-way ANOVA with Kruskal-Wallis posttest with Dunn’s multiple corrections, which indicated that only the TirC ΔNT cells were significantly smaller than the Act15:GFP controls. ** = p ≤ 0.001 **D.** Microscopy of amoeba cells expressing full-length (FL) WT TirC and **E.** FL E162A TirC. WT TirC overexpression cell lines exhibited puncta while E162A TirC overexpression cells show more diffuse distribution of fluorescent signal. Images were taken 6 days post-induction. **F.** Microscopy of amoeba cells expressing inducible GFP or inducible TirC ΔNT (both WT and E162A) at 0 hours after induction and Nicotinamide (NAM) supplementation. At 0 hours post induction, the inducible GFP control cells show little to no green fluorescence indicating minimal leaky expression. No green fluorescence was observed in the TirC ΔNT overexpression cell lines. Cells exhibited amoeboid morphology and substrate adhesion characteristic of healthy cells. **G.** Microscopy of amoeba cells expressing N-terminal truncations (ΔNT) of TirC at 24 hours after induction and NAM supplementation. At 24 hours post induction, the inducible GFP control cells show diffuse green fluorescence and normal cell morphology. In contrast, cells expressing TirC ΔNT are rounded and detached (morphology and behavior associated with loss of viability and cell death) when cultured without NAM. However, this same culture grew as amoeboid, adhered cells observed in the 20mM NAM supplemented conditions. Cells expressing ΔNT E162A TirC exhibited normal morphology, adhesion, and diffuse distribution of green fluorescent signal regardless of NAM concentration in media. Exposure and contrast levels were matched across samples. Scale bar is 10 μm.

**Figure S18.**
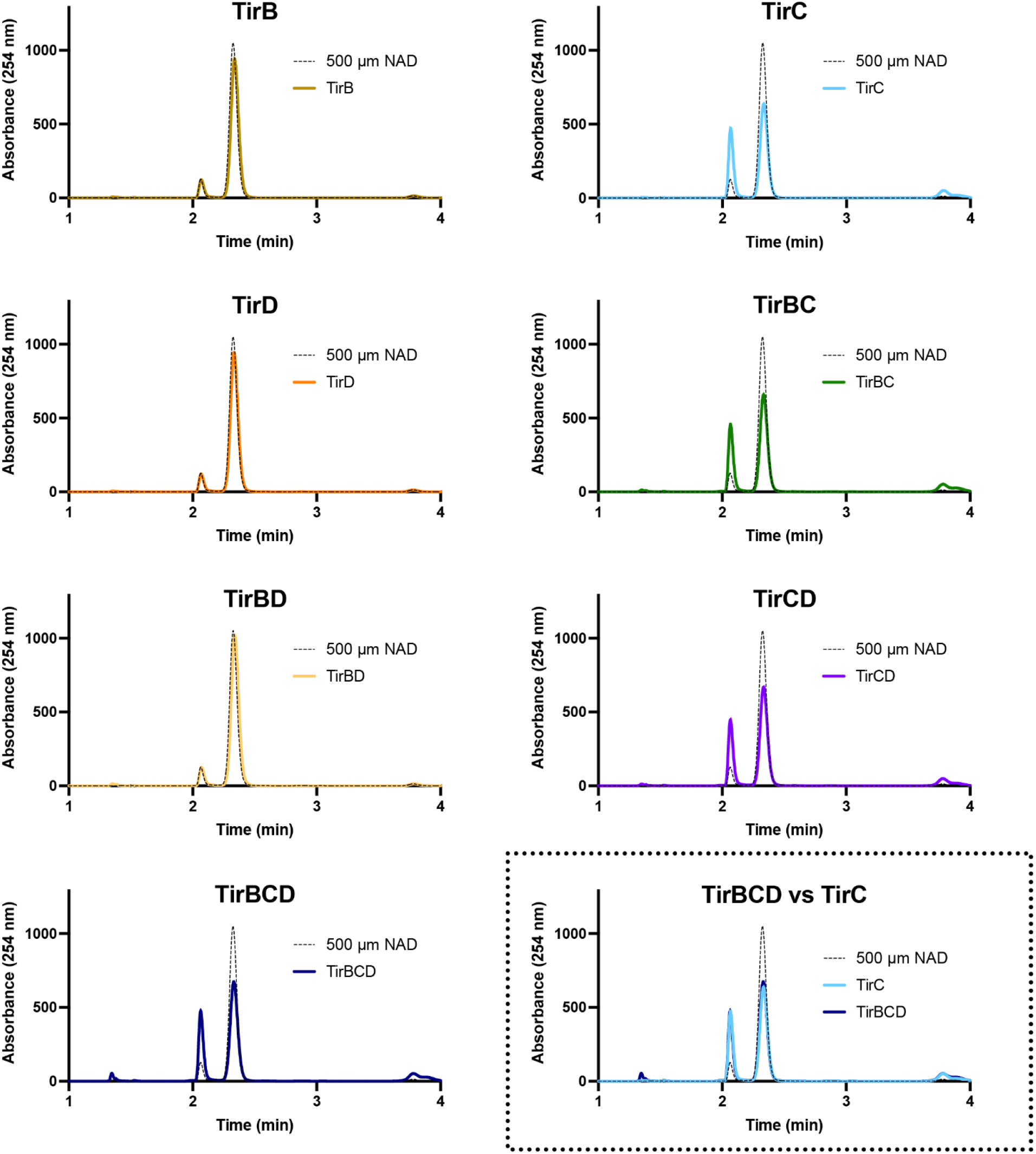
Co-incubation of TirB, C, and D proteins *in vitro* does not alter NADase activity. High-performance liquid chromatography analysis of NAD^+^ incubated with purified TirB, TirC, TirD, or combinations thereof. We observed no NAD^+^ processing from TirB or TirD, alone or with each other. In all other mixes, the observed activity was equivalent to the activity of TirC alone. The graph in the dotted box overlays the TirC-only results with those from the TirB, C, and D mix, showing that they are nearly identical.

**Figure S19.**
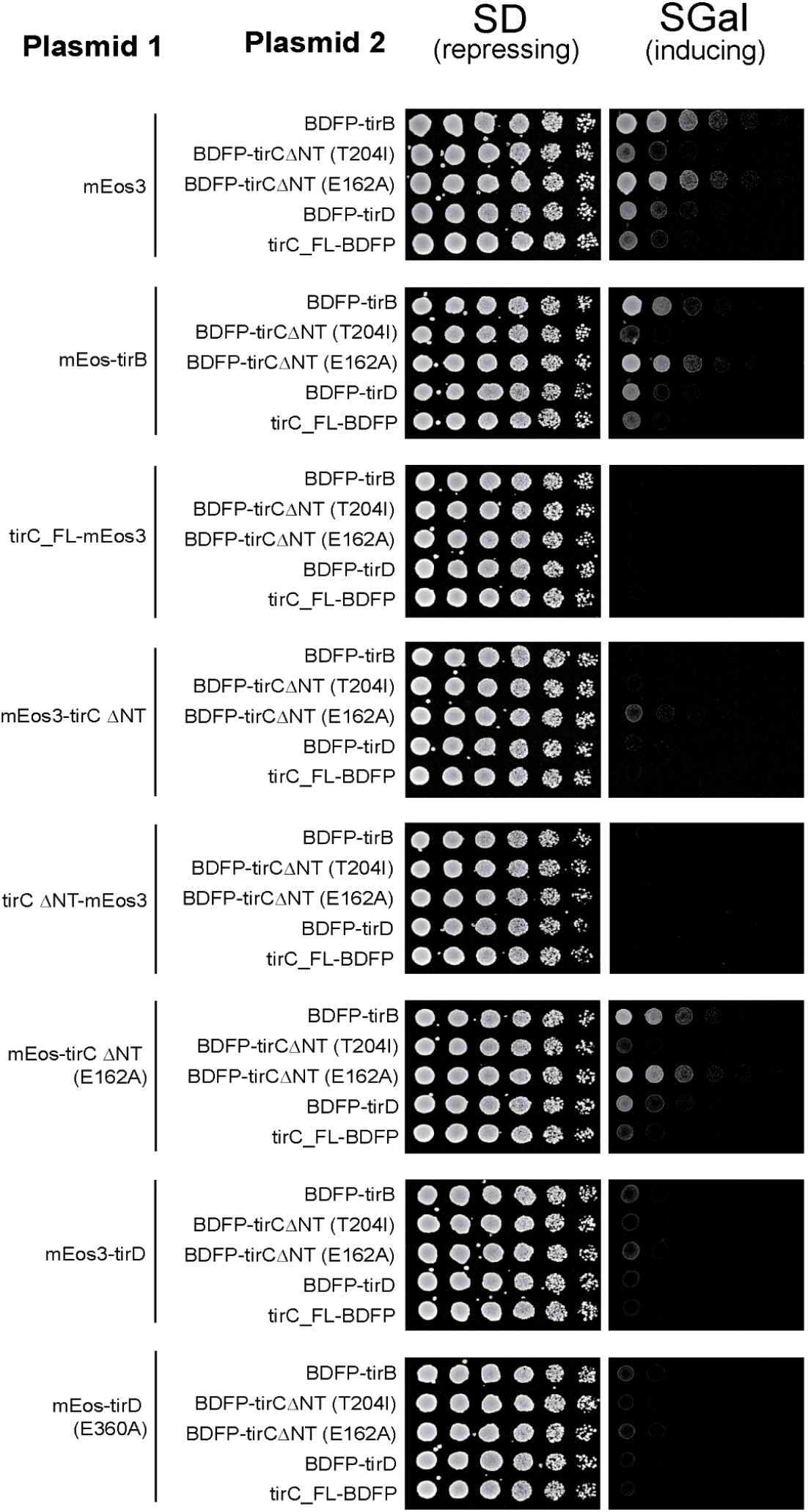
Toxicity of TirBCD co-expression in *S. cerevisiae*. Yeast cells carrying the indicated plasmid constructs were serially diluted (5-fold) and spotted onto selective media to either repress or induce co-expression of the indicated mEos- and BDFP-tagged proteins. TirC constructs were either full length (FL) or truncated to remove the N-terminal helix (ΔNT). Plates were incubated at 30°C for 3 days. Images are representative of 2 replicates of the spotting assay.

